# Testing the role of the Red Queen and Court Jester as drivers of the macroevolution of Apollo butterflies

**DOI:** 10.1101/198960

**Authors:** Fabien L. Condamine, Jonathan Rolland, Sebastian Höhna, Felix A. H. Sperling, Isabel Sanmartín

## Abstract

In macroevolution, the Red Queen (RQ) model posits that biodiversity dynamics depend mainly on species-intrinsic biotic factors such as interactions among species or life-history traits, while the Court Jester (CJ) model states that extrinsic environmental abiotic factors have a stronger role. Until recently, a lack of relevant methodological approaches has prevented the unraveling of contributions from these two types of factors to the evolutionary history of a lineage. Here we take advantage of the rapid development of new macroevolution models that tie diversification rates to changes in paleoenvironmental (extrinsic) and/or biotic (intrinsic) factors. We inferred a robust and fully-sampled species-level phylogeny, as well as divergence times and ancestral geographic ranges, and related these to the radiation of Apollo butterflies (Parnassiinae) using both extant (molecular) and extinct (fossil/morphological) evidence. We tested whether their diversification dynamics are better explained by a RQ or CJ hypothesis, by assessing whether speciation and extinction were mediated by diversity-dependence (niche filling) and clade-dependent host-plant association (RQ) or by large-scale continuous changes in extrinsic factors such as climate or geology (CJ). For the RQ hypothesis, we found significant differences in speciation rates associated with different host-plants but detected no sign of diversity-dependence. For CJ, the role of Himalayan-Tibetan building was substantial for biogeography but not a driver of high speciation, while positive dependence between warm climate and speciation/extinction was supported by continuously varying maximum-likelihood models. We find that rather than a single factor, the joint effect of multiple factors (biogeography, species traits, environmental drivers, and mass extinction) is responsible for current diversity patterns, and that the same factor might act differently across clades, emphasizing the notion of opportunity. This study confirms the importance of the confluence of several factors rather than single explanations in modeling diversification within lineages.

## Introduction

Evolutionary biologists have long endeavored to determine which factors govern biodiversity dynamics, aiming to answer questions such as why some clades have diversified more than others, or why some lineages are widely distributed whereas others survive in restricted ranges (Ezard et al. 2016). Two different mechanistic macroevolutionary models have been proposed to explain the generation and maintenance of diversity. The Red Queen (RQ) model (Van Valen 1973), which stems from Darwin and Wallace, posits that diversification is driven by species-intrinsic, biotic factors such as interactions among species, species ecology, or life-history traits. The Court Jester (CJ) model, which builds on paleontological evidence (Barnosky 2001), argues that diversification dynamics result from historical abiotic forces such as abrupt changes in climate or geological tectonic events that drive speciation and extinction rates, usually acting clade-wide across lineages.

The CJ and RQ models are generally considered the two extremes of a continuum, operating over different geographic and temporal scales. Biotic factors such as species interactions shape ecosystems locally over short time spans, whereas abiotic factors such as climate and tectonic events shape large-scale patterns regionally and globally over millions of years (Benton 2009). However, biotic interactions can also be observed at large spatial and temporal scales (Liow et al. 2015; Silvestro et al. 2015), while Van Valen’s (1973) original RQ hypothesis is now interpreted as accepting the role of a changing environment in shaping species evolution (Voje et al. 2015).

Although both abiotic (environmental) and biotic (species-intrinsic) drivers are recognized as fundamental for regulating biodiversity (Ezard et al. 2011), these two types of factors are often studied in isolation (Drummond et al. 2012a; Bouchenak-Khelladi et al. 2015; Lagomarsino et al. 2016), searching for correlations between shifts in diversification rates and the evolution of key innovations or the appearance of key opportunities (Maddison et al. 2007; Alfaro et al. 2009; Rabosky 2014; Givnish et al. 2015). However, often no single factor but a confluence of biotic and abiotic factors is responsible for the diversification rate shift (Donoghue and Sanderson, 2015), and there could be an interaction effect. For example, C_4_ grasses appeared in the Eocene but their expansion and explosive diversification started only after mid-Miocene aridification in Africa and Central Asia (Edwards et al. 2010).

Statistical assessment of the relative contributions of abiotic and biotic factors underlying diversity patterns has been made possible by the development of new probabilistic models in the field of diversification dynamics (Stadler 2013; Morlon 2014; Höhna 2015). One type of model estimates diversification rates that are clade-dependent and identifies differences in diversification rates among clades that can be explained by key innovations (Alfaro et al. 2009; Morlon et al. 2011; Rabosky et al. 2013), or by diversity-dependence and niche filling (i.e. diversification decreasing as the number of species increases, Rabosky and Lovette 2008; Etienne et al. 2012). A second type of model aims to detect statistical associations between diversification rates and changes in species traits (trait-dependent diversification models, Maddison et al. 2007; Ng and Smith 2014), or between geographic evolution and diversification, such as a change in continental connectivity allowing the colonization of a new region and a subsequent increase in allopatric speciation (Goldberg et al. 2011). A third type of model assumes continuous variation in diversification rates over time that depends on a paleoenvironmental variable and investigates whether diversification rates can be affected by abiotic environmental changes (e.g. paleotemperature, Condamine et al. 2013). Finally, episodic birth-death models search for tree-wide rate shifts that act concurrently across all lineages in a tree, for example a mass extinction event removing a fraction of lineages at a certain time in the past (Stadler 2011; Höhna 2015; May et al. 2016).

The first two types of models have been used to test RQ-like hypotheses on the effect of biotic interactions, while the other models are often used in the context of the CJ hypothesis (environmental change). Studies using a subset of these models to address both abiotic and biotic factors are becoming more frequent, but are often used at a local or regional geographic scale (Schnitzler et al. 2011; Drummond et al. 2012a; Jønsson et al. 2012; Hutter et al. 2013; Bouchenak-Khelladi et al. 2015; Lagomarsino et al. 2016), and limited temporally. No study so far has addressed the full set of models using a large (geographic) and long (temporal) scale within the same lineage. Such a study would have substantial power to provide information on questions such as why some lineages diversify and others do not, and the extent that diversification is attributable to trait evolution and/or ecological opportunity (Wagner et al. 2012). It might also shed light on the current debate about whether biodiversity is at an equilibrium bounded by ecological limits (Rabosky 2013), although it is unclear how relevant such limits are compared to other biological explanations (Moen and Morlon 2014). Alternatively, diversity might be unbounded and controlled by rare but drastic environmental changes such as climatic mass extinction events (Antonelli and Sanmartín 2011; Meredith et al. 2011; Kergoat et al. 2014; Condamine et al. 2015a; May et al. 2016).

A key to such a study would be to find a group that: *(i)* has experienced both biotic and abiotic pressures at different temporal and spatial scales; *(ii)* shows large species diversity and high specialization; and *(iii)* has good information available about its taxonomic and evolutionary history. Nearly complete taxon sampling is crucial for accurate estimation of diversification rates (Cusimano and Renner 2010; Brock et al. 2011; Höhna et al. 2011; Davis et al. 2013), whereas rich fossil evidence is important for divergence time estimation and accurate reconstruction of biogeographic history (Meseguer et al. 2015). Phytophagous insects are especially good models because they present complex biotic (trophic) interactions with their host plants (Ehrlich and Raven 1964), and many of these lineages are old enough to have experienced dramatic past climatic changes.

Here, we address the role of biotic (RQ) and abiotic (CJ) factors that drive patterns of species richness in Apollo butterflies (Papilionidae: Parnassiinae). The Parnassiinae comprise eight genera and over 70 species (Ackery 1975; Weiss 1991-2005; Kocman 2009), grouped into three tribes – Luehdorfiini, Zerynthiini, and Parnassiini (Nazari et al. 2007). Apollo butterflies occur from the Western Palearctic to the Western Nearctic (Weiss 1991-2005). Most diversity is concentrated in the Palearctic, where we find two non-monophyletic lowland-flying communities separated by the Himalaya and Tibetan Plateau (HTP, **Figure 1**): an Eastern Palearctic group formed by *Luehdorfia* (Luehdorfiini) and *Bhutanitis* and *Sericinus* (Zerynthiini), and a Western Palearctic community including *Archon* (Luehdorfiini), *Allancastria* and *Zerynthia* (Zerynthiini), and *Hypermnestra* (Parnassiini). The largest genus, *Parnassius*, is mainly distributed in mountainous regions across the Holarctic, with its highest diversity in the HTP (Weiss 1991-2005; Kocman 2009; Nazari et al. 2007; Michel et al. 2008). Compared to the species-poor genera *Allancastria*, *Archon*, *Bhutanitis*, *Hypermnestra*, *Luehdorfia* and *Sericinus* (each with five species maximum), the 50+ species of *Parnassius* have been suggested as an example of rapid diversification. This unbalanced species richness, together with their disjunct pattern of distribution in the Himalayan region, suggests that the HTP orogeny might have been an important driver in the biogeographic and diversification history of Parnassiinae.

**Figure 1.**
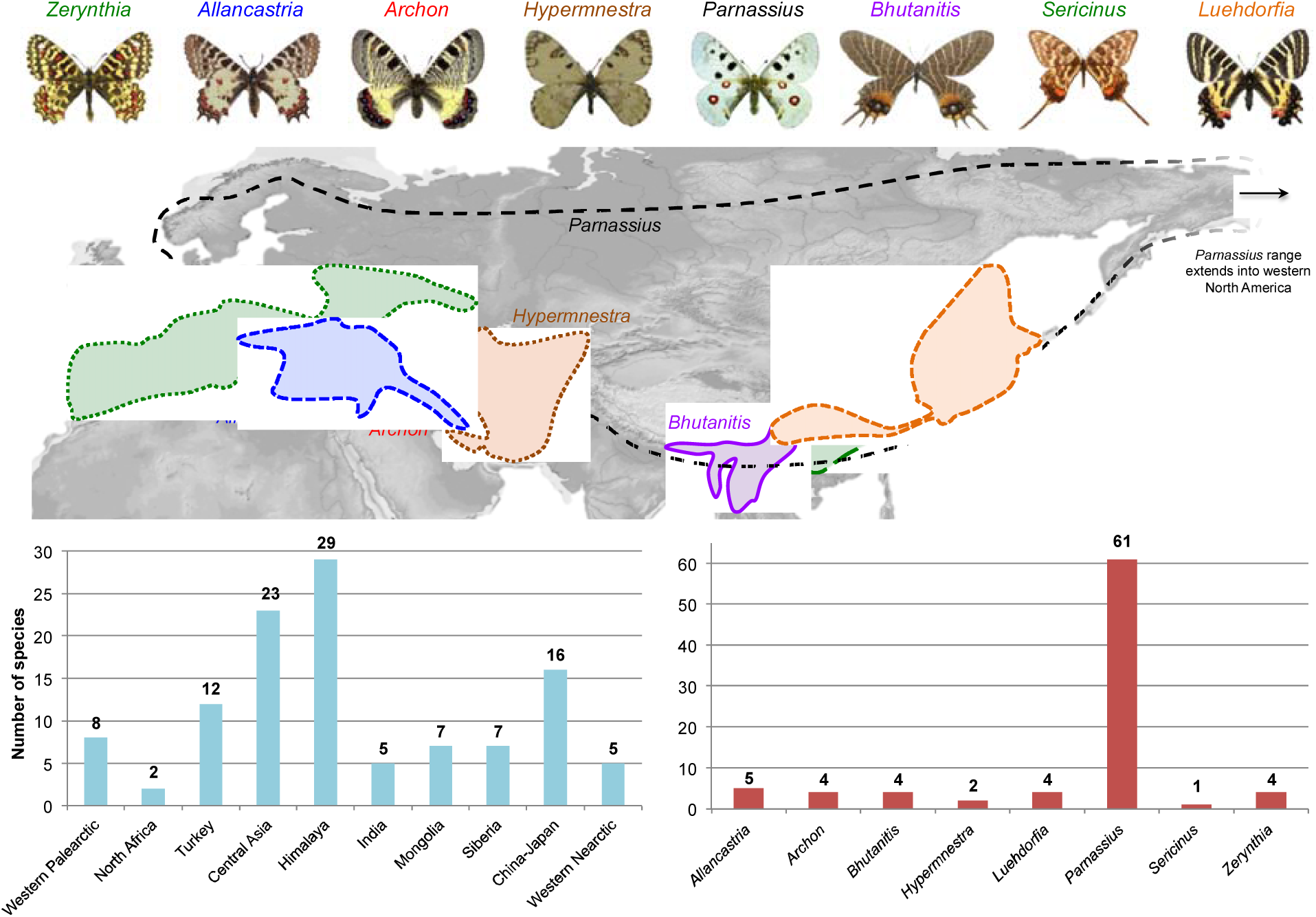
Distribution and species richness patterns of Apollo butterflies. Map shows the current separation of west and east communities of Parnassiinae, divided by the Himalayan-Tibetan Plateau. Lower portion of figure compares the current species diversity of each biogeographical unit (which may be non-monophyletic) or genus. Images of Parnassiinae (also in subsequent figures) are created by Fabien Condamine.

Parnassiinae are also remarkable for their (nearly) strict but diverse host-plant specialization (Ehrlich and Raven 1964). Tribes Luehdorfiini and Zerynthiini feed on Aristolochiaceae, the ancestral host of Parnassiinae (Condamine et al. 2012). Within Parnassiini, *Hypermnestra* feeds on Zygophyllaceae, while *Parnassius* has experienced two host-plant shifts, toward Crassulaceae+Saxifragaceae and Papaveraceae (Michel et al. 2008; Condamine et al. 2012), and is regarded as an adaptive radiation (Rebourg et al. 2006). Finally, Apollo butterflies constitute a model to understand the role of past environmental change in lineage diversification. Phylogeographic studies pinpointed the effect of Pleistocene glaciations on population-level dynamics (e.g. Keyghobadi et al. 2005; Gratton et al. 2008; Schoville and Roderick 2009; Dapporto 2010; Todisco et al. 2010, 2012; Zinetti et al. 2013). Time-calibrated phylogenies support an origin of the subfamily in the Early Cenozoic (Condamine et al. 2012), which means that the lineage has experienced the dramatic cooling and warming events of the last 50 Ma, including a drop in global temperatures at the end ofthe Eocene (Eocene-Oligocene climate transition, EOCT, Liu et al. 2009) that led to the demise of Boreotropical Holarctic vegetation (Pound et al. 2012).

In this study, we reconstruct a completely sampled phylogeny of Parnassiinae to explore the role played by abiotic (CJ) and biotic (RQ) factors, isolated or in unison, in the origin and fate of biological radiations, including paleotemperature, mountain orogeny, host specialization, range expansion and diversity dependence with niche filling. If interactions among species are the dominant drivers of evolution, diversification rates are expected to show diversity-dependent dynamics or be dependent on ecological traits. For example, one might expect diversification rates to decrease as a function of diversity or increase with ecological opportunity. Conversely, if evolution is driven mainly by changes in the physical environment, clade-wide responses due to abrupt abiotic perturbations should dominate macroevolutionary dynamics, e.g. diversification rates may shift following major climatic changes that extirpated certain lineages while favoring the radiation of others. We test these hypotheses using trait-dependent, time-dependent, environment-dependent and episodic birth-death models and compare the fit (explanatory power) of these models using maximum likelihood and Bayesian inference. Our study is the first to address the CJ and RQ hypotheses at such a large scale, and to compare the performance of these models within a single inference framework.

## Materials and Methods

### Taxon sampling and molecular dataset

The latest effort to resolve phylogenetic relationships in Parnassiinae was Condamine et al. (2012), but that study did not include all species. For our study, DNA sequences were obtained from GenBank using *Taxonomy* browser searches (full dataset is provided in the Supplementary Material that accompanies this article available on Dryad at http://dx.doi.org/10.5061/dryad.[NNNN], **Appendix 1**). Sequences of less than 250 nucleotides (possibly microsatellites) and genes sequenced in very few specimens were not included. Overall the dataset comprises >1,800 sequences, representing 69 species and five genes (four mitochondrial and one nuclear) including 730 sequences for cytochrome oxidase I (COI), 214 for NADH dehydrogenase 1 (ND1), 147 for NADH dehydrogenase 5 (ND5), 202 for rRNA 16S (16S), and 115 for elongation factor 1 alpha (EF-1α). This large molecular dataset comprises all sequences generated by previous studies on Parnassiinae, especially by F.L.C. and F.A.H.S. (Omoto et al. 2004, 2009; Katoh et al. 2005; Nazari et al. 2007; Nazari and Sperling 2007; Michel et al. 2008; Condamine et al. 2012), and includes all currently described species within the family, except for *Bhutanitis ludlowi* (Möhn 2005; Weiss 2005; Churkin 2006). The dataset also includes the majority of described subspecies or geographic races (Nazari and Sperling 2007; Gratton et al. 2008; Schoville and Roderick 2009; Dapporto 2010; Todisco et al. 2010, 2012; Zinetti et al. 2013). Additionally, sequences of nine species of swallowtails were added to the dataset as outgroups, with *Baronia brevicornis* considered as the sister group to all remaining Papilionidae, and two species for each of the four tribes included within Papilioninae, the sister subfamily of Parnassiinae (Condamine et al. 2012).

Sequences were aligned using MAFFT 7.110 (Katoh and Standley 2013) with default settings (E-INS-i algorithm). Reading frames of coding genes were checked in Mesquite 3.03 (www.mesquite.org). Datasets (species/subspecies names and GenBank accession numbers) are available on Dryad (**Appendices 1 and 2**).

### Assessment of fossil positions and calibrations

Constraints on clade ages were enforced through fossil calibrations, whose systematic position was assessed using phylogenetic analyses (Donoghue et al. 1989; Sauquet et al. 2012). Phylogenetic analyses of the dataset included both living and fossil taxa and incorporated both morphological and molecular data using a total-evidence approach (Ronquist et al. 2012a).

Swallowtail fossils are scarce, but two unambiguously belong to Parnassiinae (Nazari et al. 2007). The first is †*Thaites ruminiana* (Scudder 1875), a compression fossil from limestone in the Niveau du gypse d’Aix Formation of France (Bouches-du-Rhone, Aix-en-Provence) within the Chattian (23.03-28.1 Ma) of the late Oligocene (Rasnitsyn and Zherikhin 2002; Sohn et al. 2012). The second is †*Doritites bosniaskii* (Rebel 1898), an exoskeleton and compression fossil from Italy (Tuscany) from the Messinian (5.33-7.25 Ma, late Miocene, Sohn et al. 2012). Absolute ages of geological formations were taken from Gradstein et al. (2012).

To assess the phylogenetic positions of fossils, our morphological dataset comprised 236 characters (Hiura 1980; Nazari et al. 2007) including adult external (160) and internal (54) morphology, egg (2) and larval (10) morphology, pupal morphology (5), and characters for pigment color and chromosome number (5). Morphological characters were coded for all extant species in each genus except *Parnassius* (only eight species: one per subgenus). Yet, *Parnassius* had little impact on our analyses since there are no fossils related to this genus (Nazari et al. 2007). Fossils were only coded for adult external morphology.

Morphological and molecular data were combined to construct a total-evidence phylogeny using MrBayes 3.2.6 (Ronquist et al. 2012b). Morphological data were modeled using the Mk model (Lewis 2001). The best-fitting partitioning scheme for the molecular data was selected with PartitionFinder 1.1.1 (Lanfear et al. 2012), using the *greedy* search algorithm and the Bayesian Information Criterion (BIC). Rather than using a single substitution model per molecular partition, we sampled across the entire substitution rate model space (Huelsenbeck et al. 2004) using the reversible-jump Markov Chain Monte Carlo (rj-MCMC) option. Two independent analyses with one cold and seven heated chains each were run for 20 million generations, sampled every 2000^th^. A 50% majority rule consensus tree was built after discarding 25% samples as burn-in. The dataset used for this analysis is available on Dryad (**Appendix 3**).

### Phylogeny and estimates of divergence times

We examined whether the rate of molecular evolution evolved in a clock-like pattern using PATHd8 (Britton et al. 2007). Since the hypothesis of a strict molecular clock was rejected for 68 of the 92 nodes (*P*<0.05), we estimated divergence times using Bayesian relaxed-clock methods accounting for rate variation across lineages (Drummond et al. 2006). MCMC analyses implemented in BEAST 1.8.2 (Drummond et al. 2012b) were employed to approximate the posterior distribution of rates and divergences times and infer their credibility intervals. We set the following settings and priors: a partitioned dataset (after the best-fitting PartitionFinder scheme) was analyzed using the uncorrelated lognormal distribution (UCLD) clock model, with the mean set to a uniform prior between 0 and 1, and an exponential prior (lambda=0.333) for the standard deviation. The branching process prior was set to either a Yule or a birth-death (Gernhard 2008) process (since tree priors may impact estimates of molecular dating, Condamine et al. 2015b), using the following uniform priors: the Yule birth rate and birth-death mean growth rate ranged between 0 and 10 with a starting value at 0.1, and the birth-death relative death rate ranged between 0 and 1 (starting value=0.5).

Calibration priors employed the fossil constraints indicated above (see Results for the phylogenetic positions), using a uniform prior with the minimum age equal to the youngest age of the geological formation where the fossil was found. Additionally, we constrained the crown of Papilionidae with a uniform distribution bounded by a minimum age of 47.8 Ma (Smith et al. 2003; Gradstein et al. 2012) based on two fossils †*Praepapilio colorado* and †*P. gracilis* (Durden and Rose 1978), both from the Lutetian (mid-Eocene) of the Green River Formation (USA, Colorado). We could not place these fossils phylogenetically because of limited taxon sampling for Papilioninae. Instead, we placed these unambiguous fossils conservatively at the crown of the family as they share synapomorphies with all extant subfamilies (de Jong 2007), and have proven to be reliable calibration points for the crown group (Condamine et al. 2012). All uniform calibration priors were set with an upper bound equal to the estimated age of angiosperms (~140 Ma, *sensu* Magallón et al. 2015), which is six times older than the oldest Parnassiinae fossil. This upper age is intentionally set as ancient to allow exploration of potentially old ages for the clade. Since the fossil record of Lepidoptera is highly incomplete and biased (Sohn et al. 2015), caution is needed in using the few recent fossil calibrations.

To explore the impact of sampled diversity, tree prior, and morphological data on BEAST inferences, we ran four analyses as follows: *(i)* molecular data only and Yule model; *(ii)* molecular data only and birth-death process; *(iii)* molecular and morphological data (total-evidence analysis) and Yule model; and *(iv)* molecular and morphological data (total-evidence analysis) and birth-death process, with all other parameters set equal. The total-evidence analyses provide a completely sampled phylogeny of Parnassiinae due to the inclusion of *Bhutanitis ludlowi* (only available with morphological data).

Each MCMC analysis was run for 100 million generations, sampled every 10,000^th^esulting in 10,000 samples in the posterior distribution of which the first 2,500 samples were discarded as burn-in. All analyses were performed on the computer cluster CIPRES Science Gateway (Miller et al. 2015), using BEAGLE (Ayres et al. 2012). Convergence and performance of MCMC runs were evaluated using Tracer 1.6 (Rambaut and Drummond 2009) and the effective sample size (ESS) criterion for each parameter. A maximum-clade credibility (MCC) tree was reconstructed, with median age and 95% height posterior density (HPD). Bayes factor (BF) comparisons using stepping-stone sampling (Xie et al. 2011), which allows unbiased approximation of the marginal likelihood of Bayesian analyses (Baele et al. 2013), was used to select among competing models. We considered 2lnBF values >10 to significantly favor one model over another (Kass and Raftery 1995). The dataset and BEAST xml files generated for this study are available on Dryad (**Appendix 4**).

### Historical biogeography

Definition of biogeographic units was based on the present-day distribution of species but also informed by plate tectonics (Sanmartín et al. 2001) and alpine orogeny reconstructions (Bouilhol et al. 2013). Ten areas were defined: WN=Western North America (including the Rocky Mountains); WP=Europe (France, Spain, Pyrenees, Alps, Italy, Greece, Crete, Balkans, Scandinavia, Western Russian, and Ural Mountains); SI=Eastern Russia, Siberia, and Kamchatka; CA=Central Asia (Turkmenistan, Uzbekistan and Kazakhstan); MO=Mongolian steppes and Altai Mountains; TU=Turkey, Caucasian region, Syria, Iraq, Iranian Plateau and Zagros Mountains; HTP=Himalaya, Tibetan Plateau, and Pamir region; IN=India, Bhutan, Yunnan, and Southern China; CJ=Northern China, Korea, and Japan; AF=North Africa and Arabia.

Species ranges were defined by presence–absence in each region (**Appendix 5**). Biogeographic analyses were performed on the MCC tree (outgroups removed), using the Dispersal–Extinction–Cladogenesis model of range evolution (DEC; Ree and Smith 2008) implemented in the R-package *BioGeoBEARS* 0.2.1 (Matzke 2014). Because of concerns with statistical validity and model choice in BioGeoBEARS (Ree and Sanmartín in review), we did not use the DEC+J model (Matzke 2014). We also preferred DEC to DEC+J because the latter often infers null or extremely low extinction rates (Sanmartín and Meseguer 2016), an effect of the model favoring direct dispersal over widespread ranges, which might not be adequate for reconstructing the history of ancient groups. Biogeographic ranges larger than four areas in size were disallowed as valid biogeographic states if they were not subsets of the terminal species ranges; widespread ranges comprising areas that have never been geographically connected (Sanmartín et al. 2001) were also removed. We constructed a time-stratified model (**Appendix 6** on Dryad) with four time slices that specified constraints on area connections and spanned the Parnassiinae evolutionary history; each interval represents a geological period bounded by major changes in tectonic and climatic conditions thought to have affected the distribution of these butterflies.

### Testing the Red Queen and Court Jester hypotheses

To understand the relative contributions of RQ versus CJ-type factors, we ran a series of macroevolutionary models using both maximum likelihood (ML) and Bayesian inference (BI). Rather than on a single tree, analyses were performed on a sample of trees from the BEAST post-burnin posterior distribution (with outgroups removed); an exception was the diversity-dependent diversification models that, due to time constraints, were run only on the MCC tree. This approach allowed us to assess the robustness of results to phylogenetic uncertainty and estimate confidence intervals for the parameter estimates. We compared the fit of each model to the data using the corrected Akaike Information Criterion (AICc) for the ML analyses, and Bayes Factor comparisons (2lnBF) for BI.

#### Diversification analyses in a Red Queen model

We first tested the hypothesis that diversity is bounded or at equilibrium, meaning that diversity expanded rapidly in early diversification, filled most niches and saturated toward the present. We explored the effect of diversity-dependence on speciation and extinction rates using the method of Etienne et al. (2012) implemented in the R-package *DDD* 2.7. We applied five different models: (*i*) speciation depends linearly on diversity without extinction, (*ii*) speciation depends linearly on diversity with extinction, (*iii*) speciation depends exponentially on diversity with extinction, (*iv*) speciation does not depend on diversity and extinction depends linearly on diversity, and (*v*) speciation does not depend on diversity and extinction depends exponentially on diversity. For each model, the initial carrying capacity was set to the current number of described species.

We also tested the effect of host-plant association on diversification rates, since previous study has pointed to a potential correlation between shifts in diversification and host-plant switches (Ehrlich and Raven 1964). We used a diversification model of the state-dependent speciation and extinction (SSE) family of models, in which extinction and speciation rates are associated with phenotypic evolution of a trait along a phylogeny (Maddison et al. 2007). In particular, we used the Multiple State Speciation Extinction model (MuSSE; FitzJohn et al. 2009) implemented in the R-package *diversitree* 0.9-7 (FitzJohn 2012), which allows for multiple character states. Larval host plant data were taken from previous work (Ehrlich and Raven 1964; Scriber 1984; Collins and Morris 1985; Tyler et al. 1994; Scriber et al. 1995). The following four character states were used: (1) Aristolochiaceae; (2) Zygophyllaceae; (3) Papaveraceae; and (4) Crassulaceae+Saxifragaceae. Data at a lower taxonomic level than plant family were not used because of the great number of multiple associations exhibited by genera that could alter the phylogenetic signal. Thirty-six different models were run to test whether speciation, extinction, or transition rates were dependent on trait evolution. Models were built with increasing complexity, starting from a model with no difference in speciation, extinction and transition rates (3 parameters) to the most complex model with different speciation, extinction, and transition rates for each character state (20 parameters). We estimated posterior density distribution with Bayesian MCMC analyses (10,000 steps) performed with the best-fitting models and resulting speciation, extinction and transition rates.

There have been concerns about the power of SSE models to infer diversification dynamics from a distribution of species traits (Davis et al. 2013; Maddison and FitzJohn 2015; Rabosky and Goldberg 2015). First, SSE models tend to have a high type I error bias related to the shape (topology) of the inferred tree (Rabosky and Goldberg 2015). To test whether our diversification results were potentially biased, we estimated the difference of fit (ΔAIC) between the best model from MuSSE and a null model (corresponding to the model with no difference in speciation, extinction and transition rates between the states of a character) and compared this with the difference between the same models as estimated from simulated datasets. 1000 sets of traits were generated on the Parnassiinae phylogeny using the *sim.history* function from the R-package *phytools* (Revell 2012). To keep the phylogenetic signal in the trait, transition rates between states were simulated using values obtained from the null model fitted on the observed phylogeny.

Second, there is concern whether SSE models are uncovering actual drivers of diversification, or whether they are simply pointing to more complex patterns involving unmeasured and co-distributed factors in the phylogeny (Beaulieu and O’Meara 2016). A model with an additional hidden character may alleviate this issue. We applied the Hidden SSE (HiSSE) model to specifically account for the presence of unmeasured factors that could impact diversification rates estimated for the states of any observed trait. The analyses were performed in the Bayesian software RevBayes (Höhna et al. 2016a).

To provide an independent assessment of the relationship between diversification rates and host specificity, we used models that allow diversification rates to vary among clades. BAMM 2.5 (Rabosky et al. 2013, 2014a) was used to explore for differential diversification dynamic regimes among clades differing in their host-plant feeding. BAMM analyses were run for 10 million generations, sampling every 10,000^th^ and with four different values of the compound Poisson prior (CPP) to ensure the posterior is independent of the prior (Moore et al. 2016). Mixing and convergence among runs (ESS>200 after 15% burn-in) were assessed with the R-package *BAMMtools* 2.1 (Rabosky et al. 2014b). BAMM has been criticized for incorrectly modeling rate-shifts on extinct lineages, i.e. unobserved (extinct or unsampled) lineages inherit the ancestral diversification process and cannot experience subsequent diversification-rate shifts (Moore et al. 2016, but see Rabosky et al. 2017). To solve this, we used here a novel approach implemented in RevBayes that models rate shifts consistently on extinct lineages by using the SSE framework (Moore et al. 2016; Höhna et al. in prep.). Although there is no information of rate shifts for unobserved/extinct lineages in a phylogeny including extant species only, these types of events must be accounted for in computing the likelihood. The number of rate categories is fixed in the analysis but RevBayes allows any number to be specified, thus allowing direct comparison of different macroevolutionary regimes.

#### Diversification analyses in a Court Jester model

We evaluated the impact of abiotic factors such as abrupt changes in tectonic or climatic settings or mass extinction events (e.g. the EOCT at 33.9 Ma) using episodic birth-death models implemented in the R-packages *TreePar* 3.3 (Stadler 2011) and *TESS* 2.1 (CoMET model, Höhna et al. 2016b; May et al. 2016). These models allow detection of discrete changes in speciation and extinction rates concurrently affecting all lineages in a tree. Both approaches estimate changes in diversification rates at discrete points in time, but can also infer mass extinction events, modeled as sampling events in the past in which the extant diversity is reduced by a fraction. Speciation and extinction rates can change at those points but remain constant within time intervals. The underlying likelihood function is identical in the two methods (Stadler 2011; Höhna 2015), but TreePar and TESS differ in the inference framework (ML vs. BI) and the method used for model comparison (AICc vs. BF). In addition, TESS uses independent CPPs to simultaneously detect mass extinction events and discrete changes in speciation and extinction rates, while TreePar estimates the magnitude and timing of speciation and extinction changes independently to the occurrence of mass extinctions (i.e. the three parameters cannot be estimated simultaneously due to parameter identifiability issues, Stadler, 2011). To compare inferences between CoMET and TreePar, we performed two independent TreePar analyses allowing and disallowing mass extinction events (argument ME=TRUE/FALSE). We compared models with 0, 1 or 2 rate shifts/mass extinction events in TreePar using the AICc, while BF comparisons were used to assess model fit between models with varying number and time of changes in speciation/extinction rates and mass extinctions in TESS. Finally, we used an implementation of CoMET in RevBayes (Höhna, unpublished), an episodic birth-death model in which speciation and extinction are allowed to vary at discrete points in time, but are autocorrelated (instead of independent/uncorrelated as in CoMET) across time intervals; a Brownian model with rates in the next time interval centered around the rates in the current time interval was used. Also, the number of diversification rate shifts was set *a priori* as in population skyline models (Strimmer and Pybus 2001), while in CoMET this is modeled through a CPP prior.

To examine the influence of elevational distribution (lowland, mountain, or both) on speciation and extinction rates, we used the trait-dependent diversification model GeoSSE (Geographic State Speciation and Extinction, Goldberg et al. 2011). Geographic characters require a third widespread state, because, unlike morphological traits, ancestors can be present in more than one state (area). This requires the modeling of cladogenetic state change, the probability associated with the division of an ancestral range between the two descendants, alongside anagenetic state change (i.e. the probability of character change along branches). Species were coded by their elevation zone, using data from literature, museum records, and field observations. We evaluated 12 models of increasing complexity to test the relationship between elevational distributions and diversification. As in MuSSE above, we evaluated the performance of GeoSSE using simulation tests. Trait simulation was more complex here because there is no direct transition between states A and B. Instead, these transitions involve range expansion to an intermediate widespread state AB (dAB, dBA) followed by local extinction (xA, xB). Following Goldberg et al. (2011), we considered transition rates between A to B as null because they involve more than one instantaneous event; the transition rate from A to AB was coded as range expansion dA, from B to AB as dB; transitions from AB to A and from AB to B (“extirpation rates“) were coded as xB and xA, respectively. For our 1000 simulations, we used the transition rates of the model with equal speciation rates (sA=sB=sAB), but different extinction (xA≠xB) and transition rates (dA≠dB).

Finally, we tested the impact of paleoenvironmental variables on diversification rates, using a birth-death likelihood method in which rates may change continuously with an environmental variable, itself varying through time (Condamine et al. 2013). We tested four models: *(i)* a constant-rate birth-death model in which diversification is not associated to the environmental variable; *(ii)* speciation rate varies according to environment and extinction rate does not vary; *(iii)* speciation rate does not vary and extinction rate varies according to environment; and *(iv)* both speciation and extinction rates vary according to environment. We also tested the corresponding models in which speciation and/or extinction are allowed to vary with time, but independently from the environmental variable (time-dependent birth-death models, Morlon et al. 2011). When rates varied with the environment (E), we assumed exponential variation, such that λ(E)=λ_0_×^αE^ and µ(E)=µ_0_× e^βE^, in which and are the speciation and extinction rates for a given environmental variable for which the value is 0, and α and β are the rates of change according to the environment. Positive values for α or β mean a positive effect of the environment on speciation or extinction (and conversely). As environmental variables (**Figure 2**), we used paleotemperature (data retrieved from Zachos et al. 2008) and paleo-elevation of the HTP (a proxy for mountain building), with data compiled from existing data in the literature available on Dryad (**Appendix 7**). The R-package *pspline* was used to build environmental vectors from the data as input for the birth-death models.

**Figure 2.**
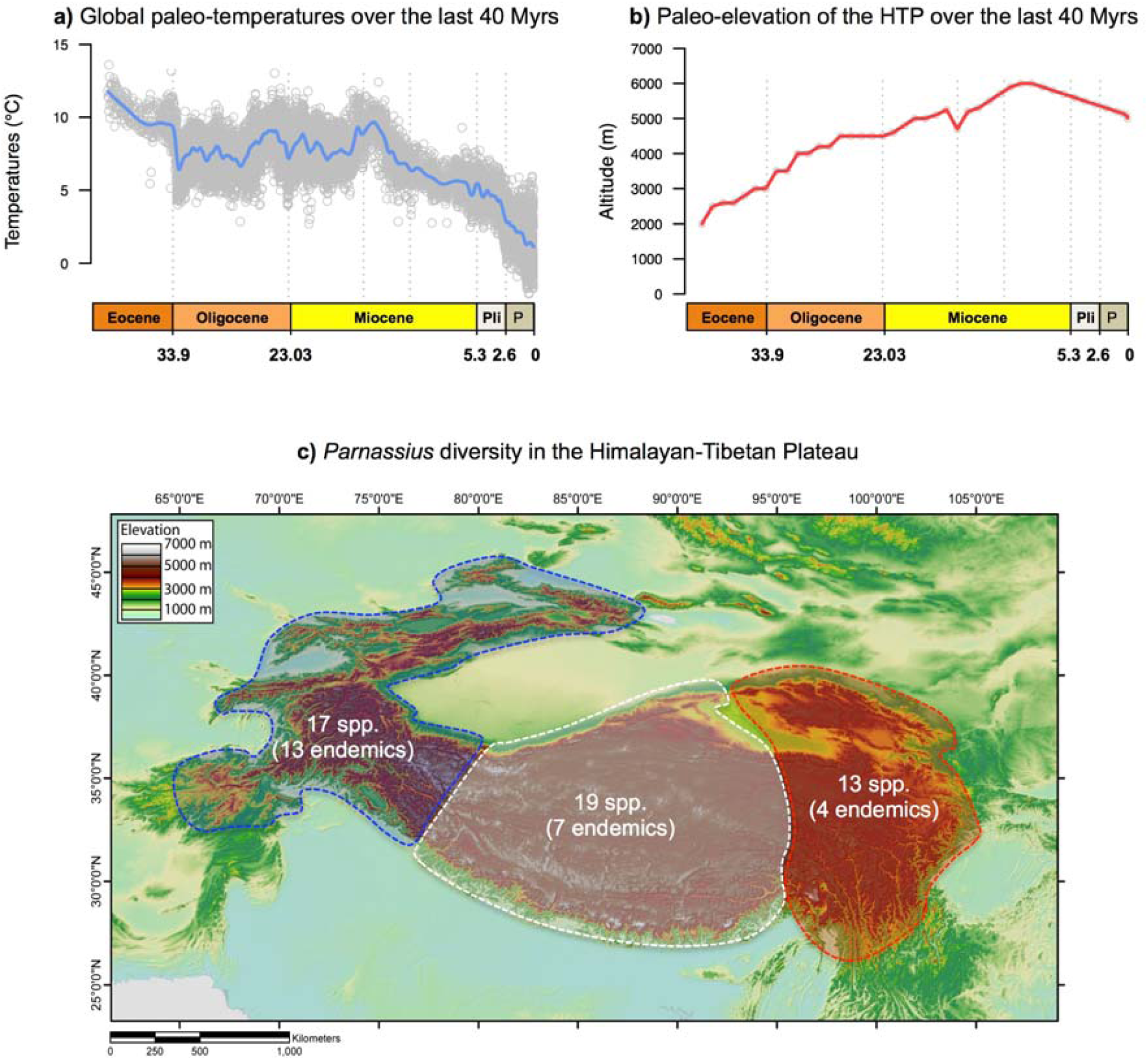
Environmental changes over the last 40 million years. a) Trends in global climate change estimated from relative proportions of different oxygen isotopes (Zachos et al. 2008), and b) trends in Himalayan-Tibetan Plateau (HTP) elevation for the last 40 Myrs, compiled from data from a literature survey; our compilation shows that the HTP was as high as 3000 m in the Eocene, congruent with recent reviews (Renner 2016; Appendix 7). c) Map of the current elevation of the Himalayan-Tibetan Plateau with the species diversity of *Parnassius* in the region.

Alternatively, we used RevBayes and the CoMET framework (episodic birth-death models) to implement an analysis in which diversification rates are allowed to co-vary with, or evolve independently from, paleotemperatures and paleo-elevation across discrete time intervals (Palazzesi al. in review). A correlation coefficient, denoted β, is co-estimated with diversification rates in the model and describes the magnitude and direction of the relationship between the variable and changes in diversification rates: β<0 (negative correlation), β>0 (positive correlation), and β=0 (no correlation, in which case the model collapses to the episodic birth-death models; see **Appendix 8** on Dryad).

## Results

### Phylogeny and fossil placements

The molecular matrix comprised 4535 nucleotides and 85 extant species of Parnassiinae plus two fossil species, following recent taxonomic reviews (Frankenbach et al. 2012; Bollino and Racheli 2012) (**Appendices 1 and 2** on Dryad). PartitionFinder selected eight partitions as the best scheme for substitution models (**Appendix 9** on Dryad). Convergence among runs was supported by the low average standard deviation of split frequencies, PSRF values ≈1.0, and ESS≫1000 for many parameters. The resulting Bayesian trees were well resolved and robust: 74.4% of the nodes were recovered with PP>0.95 with MrBayes (**Figure 3**), and 83% in BEAST (70% with PP=1) when fossil taxa were removed. Phylogenetic relationships agree with previous studies (Nazari et al. 2007; Michel et al. 2008), showing Luehdorfiini+Zerynthiini as sister-tribes and sister to Parnassiini; within the latter, *Hypermnestra* is sister to the genus *Parnassius*. Most of the weak support is found within the radiation of species complexes, such as *Parnassius apollo* or in the subgenus *Kailasius*, but two basal nodes in *Parnassius* have PP<0.9.

**Figure 3.**
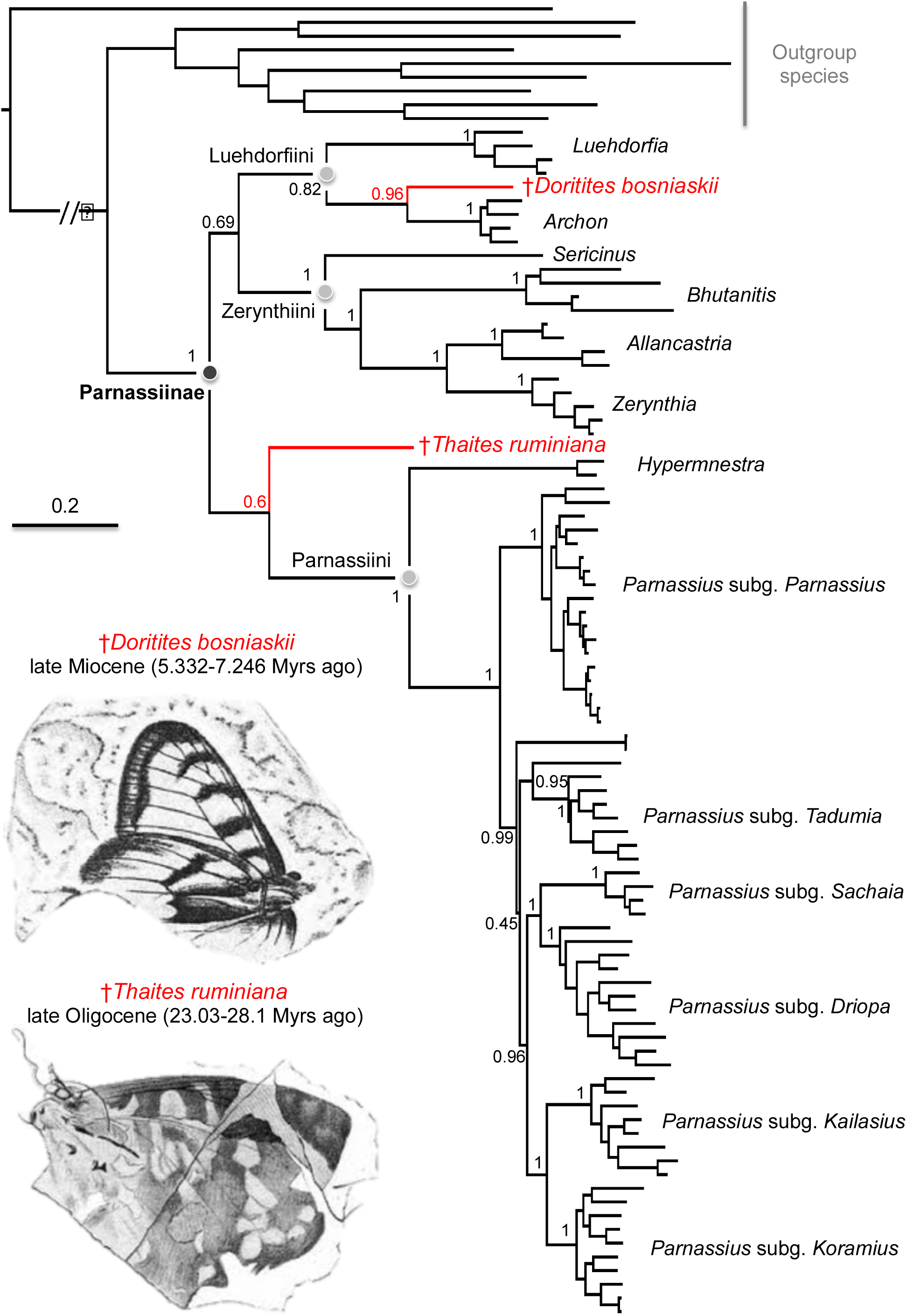
Phylogenetic relationships inferred with a Bayesian total-evidence approach combining molecular and morphological data of extant and extinct Parnassiinae. This integrative approach allowed assessment of the phylogenetic placement of two fossils, which were then used to calibrate the molecular dating analysis. Backbone nodes show posterior probability values. Nodal support values within genera or subclades are shown in Appendix 10 for alternative dating analyses including, or not, morphological data.

The total-evidence analysis allowed us to test the phylogenetic placement of the Parnassiinae fossils. †*Thaites* was often recovered as sister to Parnassiini (PP=0.6, **Figure 3**), and occasionally as sister to Luehdorfiini+Zerynthiini. This fossil was used to provide a minimum age for crown Parnassiinae, calibrated with a uniform prior bounded between 23.03 Ma (minimum) and 47.8 Ma (maximum). †*Doritites* was reconstructed as sister to *Archon* (Luehdorfiini, PP=0.96), in agreement with Carpenter (1992), who tentatively synonymized †*Doritites* with *Luehdorfia*. The crown of Luehdorfiini was thus constrained for divergence time estimation using a uniform distribution bounded between 5.33 Ma (minimum) and 47.8 Ma (maximum).

### Divergence time and biogeographic estimates

The BEAST analyses (Yule vs. birth-death, molecular vs. total-evidence) yielded almost identical estimates of divergence times with less than one million years of difference for most nodes (**Appendix 10** on Dryad). Bayes Factor comparisons were also not conclusive, with little difference in the stepping-stone marginal likelihood estimate among analyses (**Appendix 11** on Dryad). We present the results for the analysis with the birth-death prior and the total-evidence dataset (**Figure 4**) because it offered slightly better convergence and incorporating extinction into the tree prior seemed more realistic for such an old lineage (results from the other analyses are available on Dryad, **Appendix 10**).

**Figure 4.**
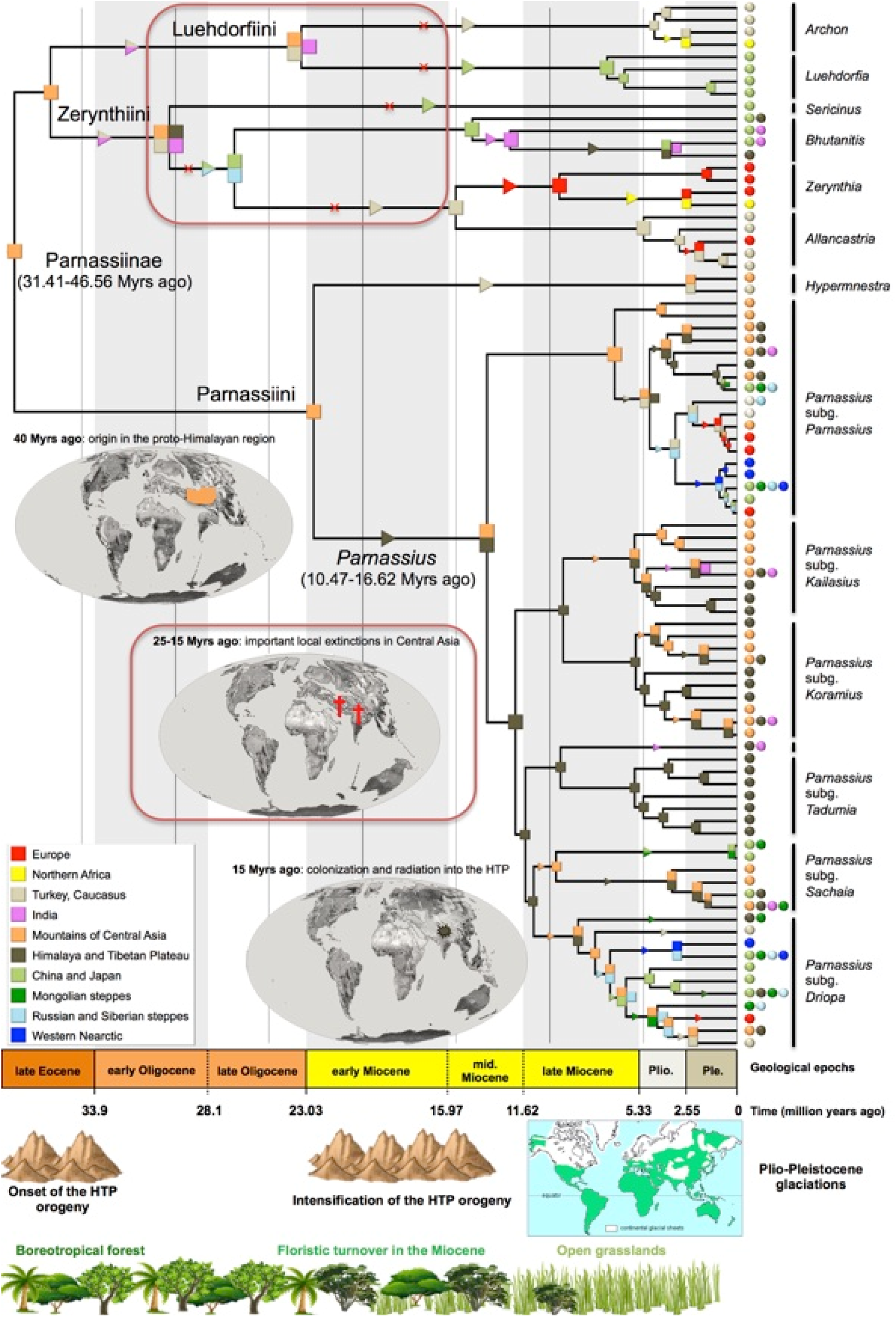
Time-calibrated phylogeny and biogeography of Apollo butterfly radiation and paleo-tectonic evolution at important periods. Colored squares on nodes indicate the most likely biogeographical estimate, as in the lower left inset (uncertainties in range estimates are in Appendix 12). Triangles indicate range expansions, and X’s denote local extinctions. A timescale spans the full evolutionary history of the group. Earth maps represent paleogeography at specific time periods. The red box highlights a notable period of local extinctions in the clade Luehdorfiini+Zerynthiini. Panels at bottom represent the main Cenozoic environmental changes occurring with the onset and rise of the Himalayan-Tibetan Plateau (HTP) or floristic turnovers. Sensitivity analyses were conducted to test the effect of adding or excluding morphological data; results are presented in Appendix 10. Plio., Pliocene; Ple., Pleistocene.

Based on this analysis and results from the DEC inference, Parnassiinae originated in the late Eocene (38.6 Ma, 95% HPD 31.4-46.6 Ma) in Central Asia (**Figure 4**). The ancestor of sister-tribes Luehdorfiini and Zerynthiini (36.7 Ma, 29.9-44.5 Ma) is also reconstructed in Central Asia, followed by independent dispersal events to India + Caucasus-Turkey and the HTP. Luehdorfiini diversified in the early Miocene at 23.3 Ma (17.4-30.2 Ma) within a broad geographic range including India, Turkey-Caucasus, and Central Asia; this was followed by extinction in Central Asia and India, and a later dispersal to China-Japan along the stem of *Luehdorfia*. Zerynthiini originated in the Oligocene (30.3 Ma, 24.1-37.2 Ma) within a region comprising India, Turkey-Caucasus, Central Asia, and the HTP, and also underwent extinction events in Central Asia and India during the Miocene. In contrast, Parnassiini first diversified within this region in the early Miocene (22.6 Ma, 17.2-28.5 Ma), followed by dispersal to Turkey-Caucasus in *Hypermnestra*. The ancestor of *Parnassius* dispersed from Central Asia to HTP around the mid-Miocene (13.4 Ma, 10.5-16.6 Ma), followed by vicariance between subgenus *Parnassius* and the ancestor of the other subgenera. Dispersal events to adjacent geographic regions and subsequent allopatric speciation are reconstructed within each subgenus, especially in *Driopa*. A more detailed account of the biogeographic history and alternative nodal reconstructions are available on Dryad (**Appendix 12**).

### Macroevolutionary dynamics

A summary of diversification rate analysis results is presented in **Table 1**.

**Table 1.** Summary of diversification analyses. Evolutionary models are divided by whether they are used to test hypotheses in favor of the Red Queen or to support the Court Jester. Abbreviations: MCC, maximum clade credibility; HTP, Himalaya and Tibetan Plateau.

| Type of birth-death | Evolutionary model | Methods used | References | Data used | Settings | Main results |
| --- | --- | --- | --- | --- | --- | --- |
| Time-dependence (rates vary continuously as a function of time and across clades) | Red Queen | BAMM<br>RevBayes | Rabosky et al. (2014)<br>Höhna et al. (2016a) | MCC tree of Parnassiinae | Bayesian model to test whether speciation rates have varied over time and/or across lineages (exponential variation) | A single shift: increase of diversification near the crown of the genus *Parnassius* |
| Diversity-dependence (rates vary as a function of the number of species) | Red Queen | DDD (*dd_ML*) | Etienne et al. (2012) | MCC tree of the clade and one of *Parnassius* (to test whether they reach their carrying capacity) | 5 models to test whether speciation declines with diversity and/or extinction increases with diversity | Both Parnassiinae and Parnassius have not reached their carrying capacity, but show some degree of diversity dependence |
| Trait-dependence (rates vary as a function of a character state for a trait) | Red Queen | MuSSE (*make.musse*)<br>RevBayes | Maddison et al. (2007)<br>FitzJohn et al. (2009)<br>Höhna et al. (2016a) | 200 posterior trees + host plant preferences (four states) | 36 models to test whether a character state impacted speciation and/or extinction and/or transition rates between host plants | Support for independent speciation rates for each host-specialist clade; no significant differences for extinction and transition rates among clades |
| Episodic birth-death model (rates vary discretely as a function of time) | Court Jester | TreePar (*bd.shifts.optim*)<br>CoMET | Stadler (2011)<br>May et al. (2016) | 200 posterior trees<br>MCC tree | 4 models to test whether diversification rates and turnover episodically changed over time | Significant diversification rate and turnover changes in the late Pliocene (3.2 Ma) and a possible mass extinction in the mid-Miocene (15 Ma) |
| Trait-dependence (rates vary as a function of a character state for a trait) | Court Jester | GeoSSE (*make.geosse*)<br>RevBayes | Maddison et al. (2007)<br>Goldberg et al. (2011)<br>Höhna et al. (2016a) | 200 posterior trees + geographical occurrence (highland, lowland, or widespread) | 12 models to test whether a character state impacted speciation and/or extinction and/or transition rates between mountains and lowlands | High speciation in mountain, medium allopatric speciation and low speciation in lowland (no effect of the trait on extinction and transition rates) |
| Environmental-dependence (rates vary as a function of a time-variable environment) | Court Jester | RPANDA (*fit_env*)<br>TESS<br>RevBayes | Condamine et al. (2013)<br>Höhna et al. (2016b)<br>Höhna (in prep.) | 200 posterior trees + past global temperatures + past altitude of the HTP | 8 models for each variable to test whether rates vary with the paleoenvironment | Speciation and extinction are positively associated with warm climate. Speciation or extinction is not significantly associated with past altitude |

#### Diversification in a Red Queen model

Diversity-dependent diversification models estimated an unlimited carrying capacity for the subfamily (**Appendix 13** on Dryad), implying that accumulating diversity did not influence diversification rates. The best-fitting DDD model was one in which extinction rate increased with diversity. For *Parnassius* the best model was a diversity-dependence process with speciation decreasing with increasing diversity. However, none of these models received significant support when compared to other models (ΔAICc<2).

MuSSE analyses on host-plant preferences selected a model in which speciation rates varied among clades feeding on different host plants; extinction and transition rates were estimated as equal (**Appendix 14** on Dryad). Zygophyllaceae feeders (*Hypermnestra*) had the lowest speciation rate, followed by Aristolochiaceae feeders (Luehdorfiini and Zerynthiini), with the highest rates for clades feeding on Papaveraceae (all *Parnassius* except subgenus *Parnassius*) and Crassulaceae+Saxifragaceae (subgenus *Parnassius*) (**Figure 5**). MCMC credibility intervals showed significant differences between the speciation rates of Papaveraceae feeders and Crassulaceae+Saxifragaceae feeders and those feeding on other host plants (http://dx.doi.org/10.5061/dryad.[NNNN], **Appendix 15**). Transition rates for host-plant switches were often estimated close to zero with strong niche conservatism in the phylogeny; once a lineage shifted to a new host it did not switch back and only rarely colonized another host plant (**Appendix 14**). We assessed the robustness of this pattern with a simulation procedure, and Bayesian implementations of HiSSE and MuSSE in RevBayes (http://dx.doi.org/10.5061/dryad.[NNNN], **Appendix 16**). All analyses supported the same diversification pattern with speciation rates being different among lineages feeding on different host plants. Our estimates were robust to known biases of SSE models, suggesting a strong effect of host plant on diversification rates of Parnassiinae.

**Figure 5.**
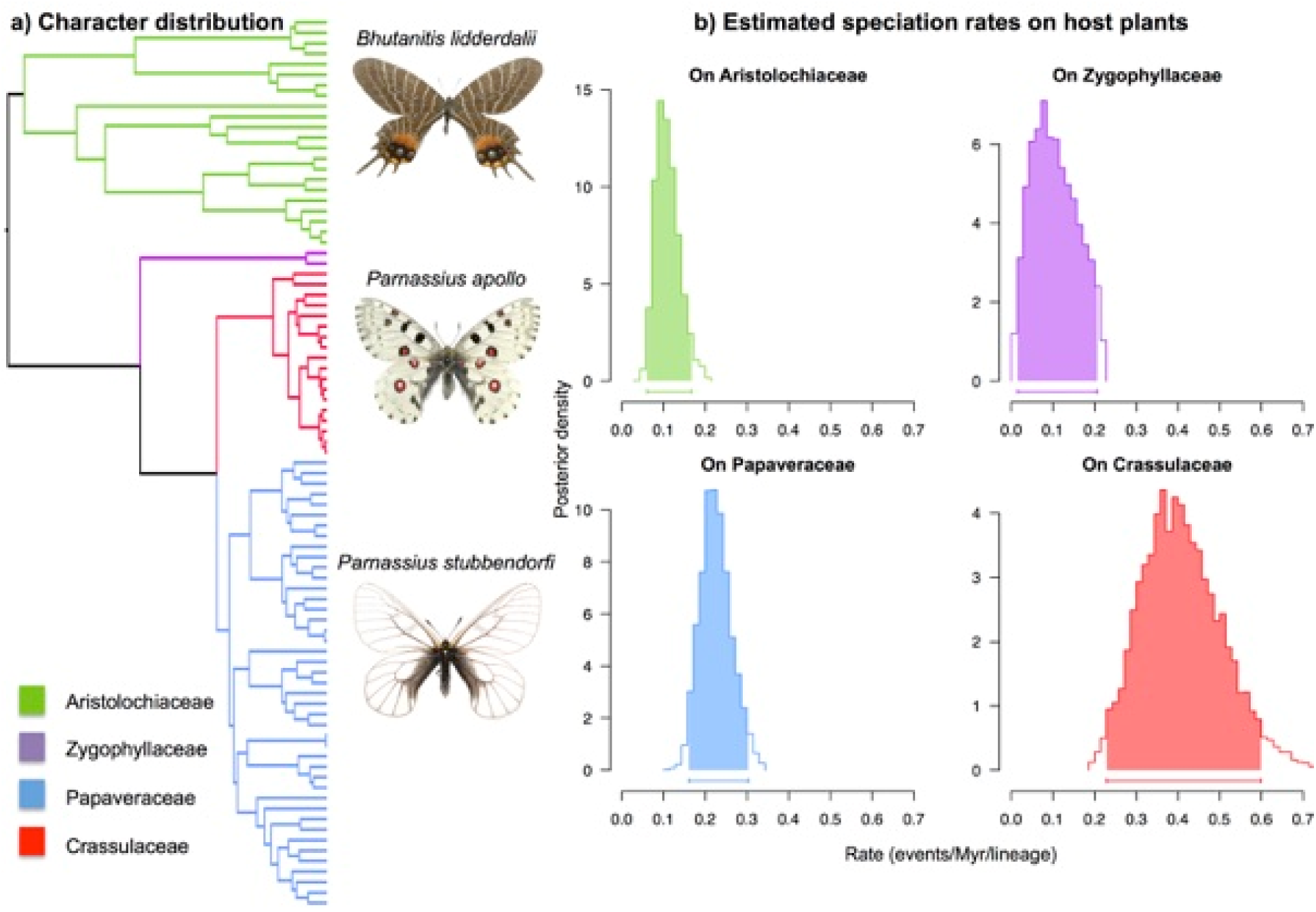
Diversification of Parnassiinae linked to their host plant. a) Phylogenetic distribution of host-plant traits (four states) analyzed with MuSSE diversification models. b) Bayesian inferences made with the best-fitting MuSSE model (see Appendix 14) showed that speciation rates vary according to the host-plant trait: *Parnassius* clades feeding on Papaveraceae and Crassulaceae+Saxifragaceae have higher speciation rates than their relatives feeding on other families; extinction and transition rates were estimated as equal across clades.

Bayesian inferences that model rate-heterogeneity across clades provided further corroboration of the SSE inferences (**Figure 6a**). Both BAMM and an alternative implementation of lineage-specific birth-death model in RevBayes supported a diversification regime in which there is at least an increase in diversification rates along the stem of *Parnassius* (and possibly a second upshift at the stem of subgenus *Parnassius*) and found higher speciation rates for host-plant feeders on Crassulaceae+Saxifragaceae and Papaveraceae than those on Aristolochiaceae or Zygophyllaceae (**Appendices 17-19** on Dryad). BAMM was insensitive to the CPP prior.

**Figure 6.**
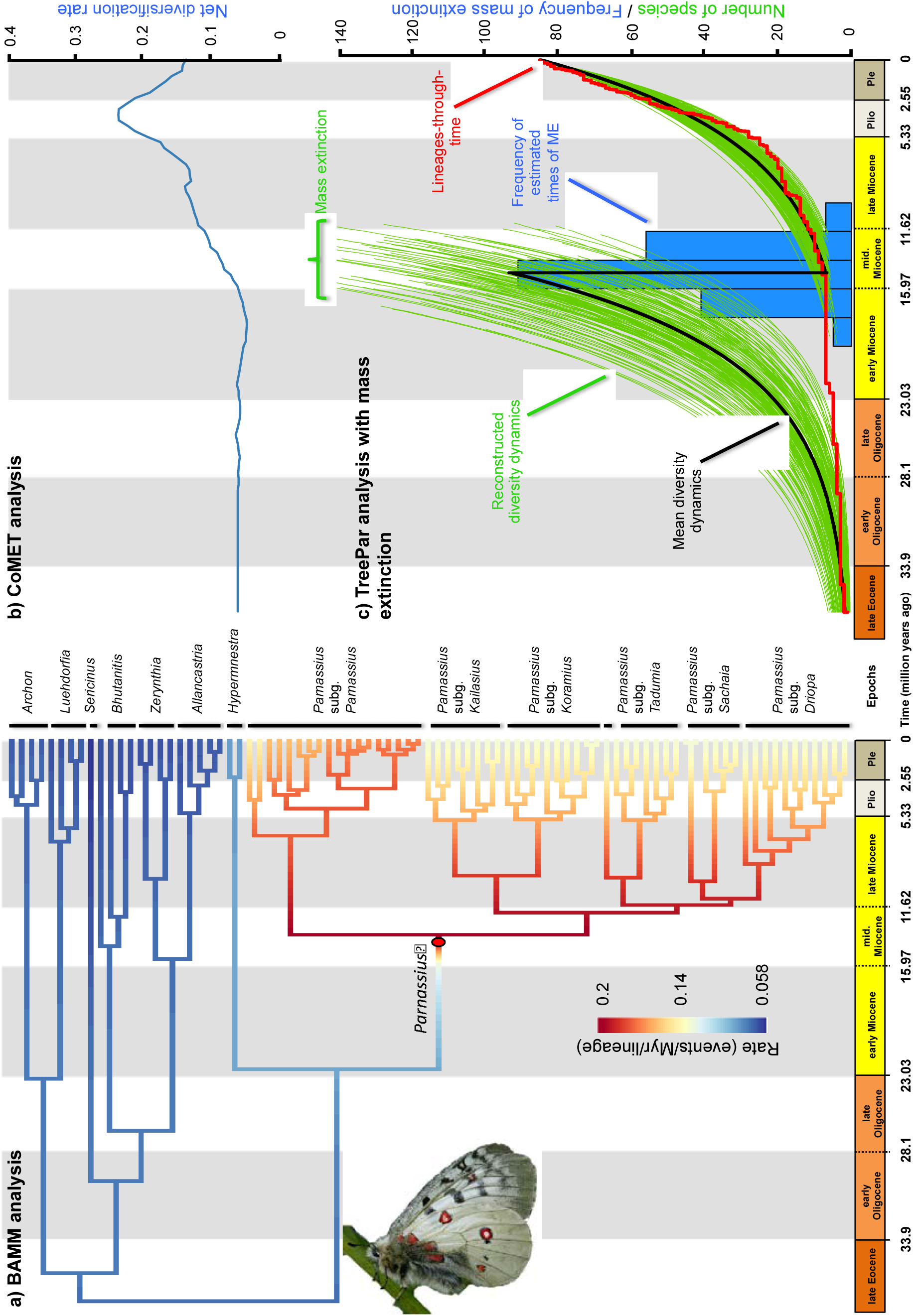
Diversification dynamics of Parnassiinae inferred with time-dependent models. a) Rate-heterogeneity-across-clades models implemented in BAMM and RevBayes identified one significant shift in diversification rates along the stem leading to *Parnassius* (represented by a red circle; see also Appendices 17-19), with a higher speciation rate in this clade than in the rest of the tree. Episodic birth-death models implemented in CoMET (b) and TreePar (c) identified a possible mass extinction event in the mid-Miocene (~15 Ma). The blue histogram shows the uncertainty in the timing of the mass extinction event based on the estimates over 200 trees with TreePar (similar estimates are obtained for CoMET; Appendix 20). The two models also estimated an increase in diversity around the early-mid Pliocene (b,c), but this was followed by a decrease in diversity towards the present time in CoMET which was not detected in TreePar. The red line shows the lineages-through-time plot. The green lines represent the reconstructed past diversity before and after the mass extinction, based on the net diversification rates and 7.5% of survival probability estimated in TreePar. Plio., Pliocene; Ple., Pleistocene.

#### Diversification in a Court Jester model

When mass extinctions are disallowed, TreePar supported a model with a single shift at 3.2 Ma (**Table 2**), with a low initial rate of diversification (r_2_=0.031) and high turnover (ε_2_=0.91), followed by 4-fold increase in diversification rate (r_1_=0.119) and a joint decrease of turnover (ε_1_=0.17). The second best model identified a second shift in the mid-Miocene (15.3 Ma) marked by an extinction period surrounded by low and negative values for the net diversification rate, and values >1 for turnover. When mass extinctions are allowed to occur, TreePar favored a significant mass extinction at 14.8 Ma with an estimated survival probability after the mass extinction of 7.5% (**Table 2**; **Figure 6c**). The CoMET model implemented in RevBayes with the autocorrelated model also identified a mass extinction event in the mid-Miocene ~15 Ma (2lnBF=6, **Appendix 20** on Dryad). There was also a significant increase in speciation rates around the Pliocene, followed by a decrease in speciation towards the present (**Figure 6b**); similar results were obtained with the TESS implementation of CoMET (**Appendix 20**).

**Table 2.** Results from episodic diversification analyses in TreePar with or without mass extinction. When mass extinction is disallowed, the best-supported model has a single rate-shift, as determined by the lowest ΔAICc. When mass extinction is allowed, the best model also has a single mass-extinction event. Adding more diversification shifts or mass extinctions did not significantly improve the likelihood of the models. Abbreviations: BD, birth-death; ME, mass extinction; NP, number of parameters; logL, log-likelihood; AICc, corrected Akaike Information Criterion; ΔAICc, the difference in AICc between the model with the lowest AICc and the others. Parameter estimates: r, net diversification rate (speciation minus extinction); ε, turnover (extinction over speciation); st, shift time; sp, survival probability at a mass extinction event. ‘r_1_’ denotes the diversification rate and ‘ε_1_’ is the turnover, both inferred between Present and the shift time 1 (‘st1’).

| Mass extinction events disallowed |  |  |  |  | Mass extinction events allowed |  |  |  |
| --- | --- | --- | --- | --- | --- | --- | --- | --- |
| Models | BD with no shift time | BD with 1 shift time | BD with 2 shift times | BD with 3 shift times | Models | BD with no ME | BD with 1 ME | BD with 2 MEs |
| NP | 2 | 5 | 8 | 11 | NP | 2 | 4 | 6 |
| logL | -182.840 | **-175.099** | -172.910 | -170.504 | logL | -240.789 | **-235.792** | -234.082 |
| | $\pm 0.5640$ | $\pm 0.5727$ | $\pm 0.5716$ | $\pm 0.5778$ | | $\pm 0.5634$ | $\pm 0.5667$ | $\pm 0.5760$ |
| AICc | 369.827 | **360.959** | 363.714 | 366.624 | AICc | 485.726 | **482.345** | 486.058 |
| | $\pm 1.1281$ | $\pm 1.1454$ | $\pm 1.1432$ | $\pm 1.1555$ | | $\pm 1.1267$ | $\pm 1.1334$ | $\pm 1.1520$ |
| ΔAIC | 8.868 | **0** | 2.755 | 5.665 | ΔAIC | | | |
| r1 | 0.0934 | **0.1188** | 0.0915 | 0.0711 | r | 0.0930 | **0.1672** | 0.1859 |
| | $\pm 0.0008$ | $\pm 0.0057$ | $\pm 0.0067$ | $\pm 0.0105$ | | $\pm 0.0008$ | $\pm 0.0012$ | $\pm 0.0014$ |
| ε1 | 0.5374 | **0.1699** | 0.3109 | 0.5288 | $\epsilon$ | 0.5363 | **0.0279** | 0.0025 |
| | $\pm 0.0047$ | $\pm 0.0355$ | $\pm 0.0411$ | $\pm 0.1224$ | | $\pm 0.0042$ | $\pm 0.0038$ | $\pm 0.0010$ |
| st1 | - | **3.21** | 2.81 | 2.56 | st1 | - | **14.82** | 7.37 |
| | | $\pm 0.0573$ | $\pm 0.0693$ | $\pm 0.0788$ | | | $\pm 0.12$ | $\pm 0.17$ |
| r2 | - | **0.0312** | 0.0254 | 0.0031 | sp1 | - | **0.0749** | 0.466 |
| | | $\pm 0.0007$ | $\pm 0.0166$ | $\pm 0.0295$ | | | $\pm 0.0021$ | $\pm 0.0144$ |
| ε2 | - | **0.9142** | 0.8319 | 0.9179 | st2 | - | - | 16.08 |
| | | $\pm 0.0052$ | $\pm 0.0529$ | $\pm 0.0795$ | | | | $\pm 0.29$ |
| st2 | - | - | 15.29 | 10.47 | sp2 | - | - | 0.0759 |
| | | | $\pm 0.8932$ | $\pm 0.6951$ | | | | $\pm 0.0028$ |
| r3 | - | - | -0.3498 | 0.0302 |  |  |  |  |
| | | | $\pm 0.1026$ | $\pm 0.0190$ | | | | |
| ε3 | - | - | 0.7616 | 0.8461 | | | | |
| | | | $\pm 0.0380$ | $\pm 0.0506$ | | | | |
| st3 | - | - | - | 20.29 |  |  |  |  |
| | | | | $\pm 0.9277$ | | | | |
| r4 | - | - | - | -0.9929 |  |  |  |  |
| | | | | $\pm 0.1753$ | | | | |
| ε4 | - | - | - | 0.8571 | | | | |
| | | | | $\pm 0.0403$ | | | | |

GeoSSE analyses selected a best-fitting model with equal dispersal and extinction rates between altitudinal regions (habitats), but higher speciation rates for highland regions compared to lowland regions (**Appendix 14** on Dryad). Allopatric speciation (for lowland/highland ancestors) was inferred to occur at a higher rate than within-lowland speciation, but at a lower rate than within-highland speciation; these differences remained significant in the Bayesian MCMC analyses, as the credibility interval did not overlap with zero (**Figure 7**). Nevertheless, the difference in speciation rates between higher and lower altitudes was not larger than expected by chance in our simulation tests (**Appendix 16** on Dryad), so we cannot reject the observed effect of altitude on speciation being affected by the shape of the phylogeny.

**Figure 7.**
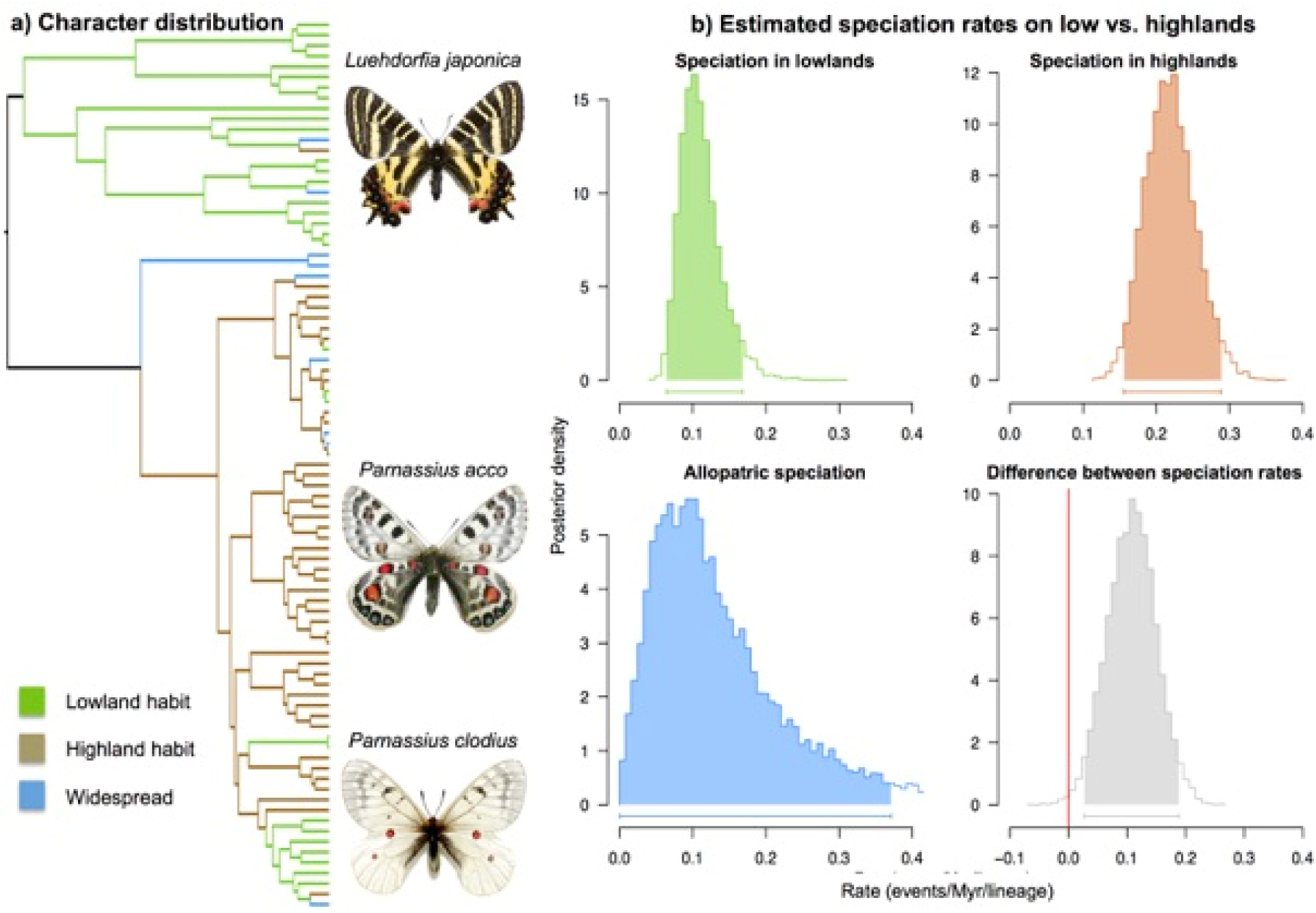
Inference of geographic mode of speciation between mountain and lowland species. a) The phylogenetic distribution of geographic traits (lowland, mountain or ‘widespread’) analyzed with GeoSSE models of diversification. b) Bayesian inferences made with the best-fitting GeoSSE model (selected over a series of 12 diversification models) showing that only speciation varies according to the geographic trait. The results indicate that the mountain-dwelling Parnassiinae species have significantly higher speciation rates than their counterparts. The allopatric (or biome divergence) speciation rate is also estimated to be elevated, congruent with our biogeographic estimates.

The ML-based paleoenvironment-dependent model implemented in RPANDA indicated positive dependence between diversification rates and past temperature (**Table 3**). In the best model, both speciation and extinction rates increased exponentially with temperature, with a general decrease in diversification rates toward the present, combined with periods of high turnover rates in the late Oligocene and mid-Miocene (**Figure 8**). The best paleo-elevation model had speciation increasing with increasing altitude, whereas extinction remained constant (and not affected by elevation fluctuations), but this model does not fit better than constant-rate or temperature-dependent models (**Table 3**). In contrast, the Bayesian paleoenvironment-dependent model implemented in RevBayes indicated that both past temperature and elevation did not impact the diversification of Parnassiinae, since the correlation parameters α and β had credibility intervals that were not significantly different from zero (**Appendix 21** on Dryad). Moreover, estimates with this model (**Figure 8**) showed diversification dynamics that were similar to those obtained with CoMET and TreePar, but unlike those from RPANDA. These differences might be attributed to the inference framework (ML in RPANDA vs. BI in RevBayes) or to the different underlying birth-death models (continuous in RPANDA vs. episodic in RevBayes). To test this, we implemented the continuous-paleoenvironmental models (Condamine et al. 2013) in TESS. We then compared these models against a constant birth-death model, episodic (piecewise constant) birth-death models (1 to 5 shifts), and time-variable models. This procedure ensured that a single statistical framework (ML and AIC) implemented in TESS was used to compare all models. Results showed that the best-fitting model was one in which speciation and extinction rates vary positively with paleotemperature, in agreement with RPANDA (**Table 4**). We therefore concluded that the different results are related not to the inference framework but to the underlying model: the use of an episodic birth-death model with autocorrelated rates across time intervals in the RevBayes model rather than the uncorrelated episodic model implemented in TESS.

**Table 3.** Results from RPANDA analyses. The model with positive dependence of speciation and extinction on temperature has the lowest AICc and lowest AICω Macroevolutionary scenarios were compared in which speciation and extinction vary through time (Time), or by paleo-elevation changes of the Himalaya and Tibetan Plateau (Alti.), and global temperature changes (Temp.). Abbreviations: NP, number of parameters; logL, log-likelihood; AICc, corrected Akaike Information Criterion; ΔAICc, the difference in AICc between the model with the lowest AICc and the others; AICω, Akaike weight. λ_0_ and µ_0_, speciation and extinction for environmental value at present; α and β, parameter controlling variation of speciation (α) and β) with paleoenvironment.

| Models | NP | logL | AICc | ΔAIC | AICω | $\lambda_0$ | $\alpha$ | $\mu_0$ | $\beta$ |
| --- | --- | --- | --- | --- | --- | --- | --- | --- | --- |
| Constant birth-death | 2 | -240.336<br>$\pm 0.620$ | 484.819<br>$\pm 1.240$ | 5.186 | 0.044 | 0.2048<br>$\pm 0.0018$ | - | 0.1123<br>$\pm 0.0017$ | - |
| $\lambda_{\text{Time}}$ and $\mu_{\text{constant}}$ | 3 | -238.427<br>$\pm 0.624$ | 483.151<br>$\pm 1.249$ | 3.518 | 0.101 | 0.1962<br>$\pm 0.0016$ | -0.0407<br>$\pm 0.0004$ | 5.00E-08<br>$\pm 5.00$E-08 | - |
| $\lambda_{\text{constant}}$ and $\mu_{\text{Time}}$ | 3 | -239.089<br>$\pm 0.621$ | 484.474<br>$\pm 1.242$ | 4.841 | 0.052 | 0.1914<br>$\pm 0.0016$ | - | 0.0549<br>$\pm 0.0010$ | 0.0433<br>$\pm 0.0006$ |
| $\lambda_{\text{Time}}$ and $\mu_{\text{Time}}$ | 4 | -238.427<br>$\pm 0.624$ | 485.354<br>$\pm 1.249$ | 5.721 | 0.034 | 0.1962<br>$\pm 0.0016$ | -0.0405<br>$\pm 0.00004$ | 8.535E-05<br>$\pm 3.24$E-05 | 0.0347<br>$\pm 0.0024$ |
| $\lambda_{\text{Alti.}}$ and $\mu_{\text{constant}}$ | 3 | -240.247<br>$\pm 0.572$ | 486.790<br>$\pm 1.144$ | 7.157 | 0.016 | 0.0310<br>$\pm 0.0012$ | 0.0004<br>$\pm 6.35$E-06 | 0.0794<br>$\pm 0.0021$ | - |
| $\lambda_{\text{constant}}$ and $\mu_{\text{Alti.}}$ | 3 | -241.542<br>$\pm 0.577$ | 489.379<br>$\pm 1.155$ | 9.746 | 0.004 | 0.1991<br>$\pm 0.0017$ | - | 0.5116<br>$\pm 0.0676$ | -0.00024<br>$\pm 1.35$E-05 |
| $\lambda_{\text{Alti.}}$ and $\mu_{\text{Alti.}}$ | 4 | -240.253<br>$\pm 0.582$ | 489.005<br>$\pm 1.164$ | 9.372 | 0.005 | 0.0454<br>$\pm 0.0021$ | 0.0003<br>$\pm 9.30$E-06 | 0.1016<br>$\pm 0.0058$ | -2.67E-05<br>$\pm 6.29$E-05 |
| $\lambda_{\text{Temp.}}$ and $\mu_{\text{constant}}$ | 3 | -240.605<br>$\pm 0.632$ | 487.505<br>$\pm 1.264$ | 7.872 | 0.011 | 0.2121<br>$\pm 0.0021$ | -0.0242<br>$\pm 0.0012$ | 0.0834<br>$\pm 0.0011$ | - |
| $\lambda_{\text{constant}}$ and $\mu_{\text{Temp.}}$ | 3 | -238.058<br>$\pm 0.649$ | 482.412<br>$\pm 1.298$ | 2.779 | 0.146 | 0.1817<br>$\pm 0.0013$ | - | 0.0071<br>$\pm 0.0005$ | 0.4707<br>$\pm 0.0138$ |
| $\lambda_{\text{Temp.}}$ and $\mu_{\text{Temp.}}$ | 4 | **-235.566**<br>**$\pm 0.595$** | **479.633**<br>**$\pm 1.189$** | **0** | **0.586** | **0.0839**<br>**$\pm 0.0017$** | **0.3811**<br>**$\pm 0.0073$** | **0.0797**<br>**$\pm 0.0019$** | **0.3988**<br>**$\pm 0.0041$** |

**Table 4.** Results from the TESS analyses. a) Paleoenvironment-dependent diversification models. b) Episodic birth-death models. The model with postive dependence of speciation and extinction with temperature has the lowest AICc and highest AICω. We compared macroevolutionary scenarios in which speciation and extinction vary through time (Time), or according to paleo-elevation changes of the Himalaya and Tibetan Plateau (Alti.), according to global temperature changes (Temp.), and episodic birth-death models (Shift). Abbreviations: NP, number of parameters; logL, log-likelihood; AICc, corrected Akaike ΔAICc, the difference in AICc between the model with the lowest AICc and the others; AICω, Akaike weight. Parameter estimates: λ_0_ and μ_0_ speciation and extinction for the environmental value at present; α and β, parameter controlling the variation of the speciation (α) and extinction (β) with the paleoenvironment. λ1 and ¼ 1, speciation and extinction for the first period of time (present to first shift).

| Models | NP | logL | AICc | ΔAIC | AICω | $\lambda_0$ | $\alpha$ | $\mu_0$ | $\beta$ |
| --- | --- | --- | --- | --- | --- | --- | --- | --- | --- |
| Constant birth-death | 2 | -240.687 | 485.425 | 3.668 | 0.044 | 0.2037 | - | 0.1113 | - |
| $\lambda_{\text{Time}}$ and $\mu_{\text{constant}}$ | 3 | -238.811 | 483.775 | 2.018 | 0.101 | 0.1951 | -0.0402 | 2.01E-06 | - |
| $\lambda_{\text{constant}}$ and $\mu_{\text{Time}}$ | 3 | -239.331 | 484.816 | 3.059 | 0.060 | 0.1898 | - | 0.0515 | 0.0467 |
| $\lambda_{\text{Time}}$ and $\mu_{\text{Time}}$ | 4 | -238.802 | 485.916 | 4.159 | 0.035 | 0.1951 | -0.0398 | 0.0002 | 0.0330 |
| $\lambda_{\text{Alti.}}$ and $\mu_{\text{constant}}$ | 3 | -239.570 | 485.294 | 3.537 | 0.047 | 0.0483 | 0.0003 | 0.09450 | - |
| $\lambda_{\text{constant}}$ and $\mu_{\text{Alti.}}$ | 3 | -240.416 | 486.985 | 5.228 | 0.020 | 0.1999 | - | 0.3730 | -0.00014 |
| $\lambda_{\text{Alti.}}$ and $\mu_{\text{Alti.}}$ | 4 | -239.378 | 487.067 | 5.310 | 0.019 | 0.0472 | 0.0004 | 0.1169 | 0.0002 |
| $\lambda_{\text{Temp.}}$ and $\mu_{\text{constant}}$ | 3 | -239.570 | 485.294 | 3.537 | 0.047 | 0.0483 | 0.0003 | 0.0945 | - |
| $\lambda_{\text{constant}}$ and $\mu_{\text{Temp.}}$ | 3 | -237.826 | 481.806 | 0.049 | 0.271 | 0.1800 | - | 0.0064 | 0.5007 |
| $\lambda_{\text{Temp.}}$ and $\mu_{\text{Temp.}}$ | **4** | **-236.723** | **481.757** | **0** | **0.278** | **0.1383** | **0.1463** | **0.0458** | **0.3483** |

**b)**
| **Models** | **NP** | **logL** | **AICc** | **ΔAIC** | **AIC<sub>w</sub>** | $\lambda_1$ | $\lambda_2$ | $\lambda_3$ | $\lambda_4$ | $\lambda_5$ | $\lambda_6$ | $\mu_1$ | $\mu_2$ | $\mu_3$ | $\mu_4$ | $\mu_5$ | $\mu_6$ |
| --- | --- | --- | --- | --- | --- | --- | --- | --- | --- | --- | --- | --- | --- | --- | --- | --- | --- |
| Shift 1 | 5 | -237.008 | 484.542 | 2.785 | 0.069 | 0.052 | 0.176 | - | - | - | - | 0.004 | 0.018 | - | - | - | - |
| Shift 2 | 8 | -236.086 | 489.706 | 7.949 | 0.005 | 0.195 | 0.084 | 0.180 | - | - | - | 0.112 | 0.097 | 0.004 | - | - | - |
| Shift 3 | 11 | -236.118 | 497.379 | 15.622 | 0.0001 | 0.180 | 0.127 | 0.155 | 0.181 | - | - | 0.111 | 0.051 | 0.159 | 0.002 | - | - |
| Shift 4 | 14 | -234.251 | 501.935 | 20.178 | 0 | 0.164 | 0.159 | 0.049 | 0.161 | 0.190 | - | 0.136 | 0.062 | 0.074 | 0.058 | 0.008 | - |

**Figure 8.**
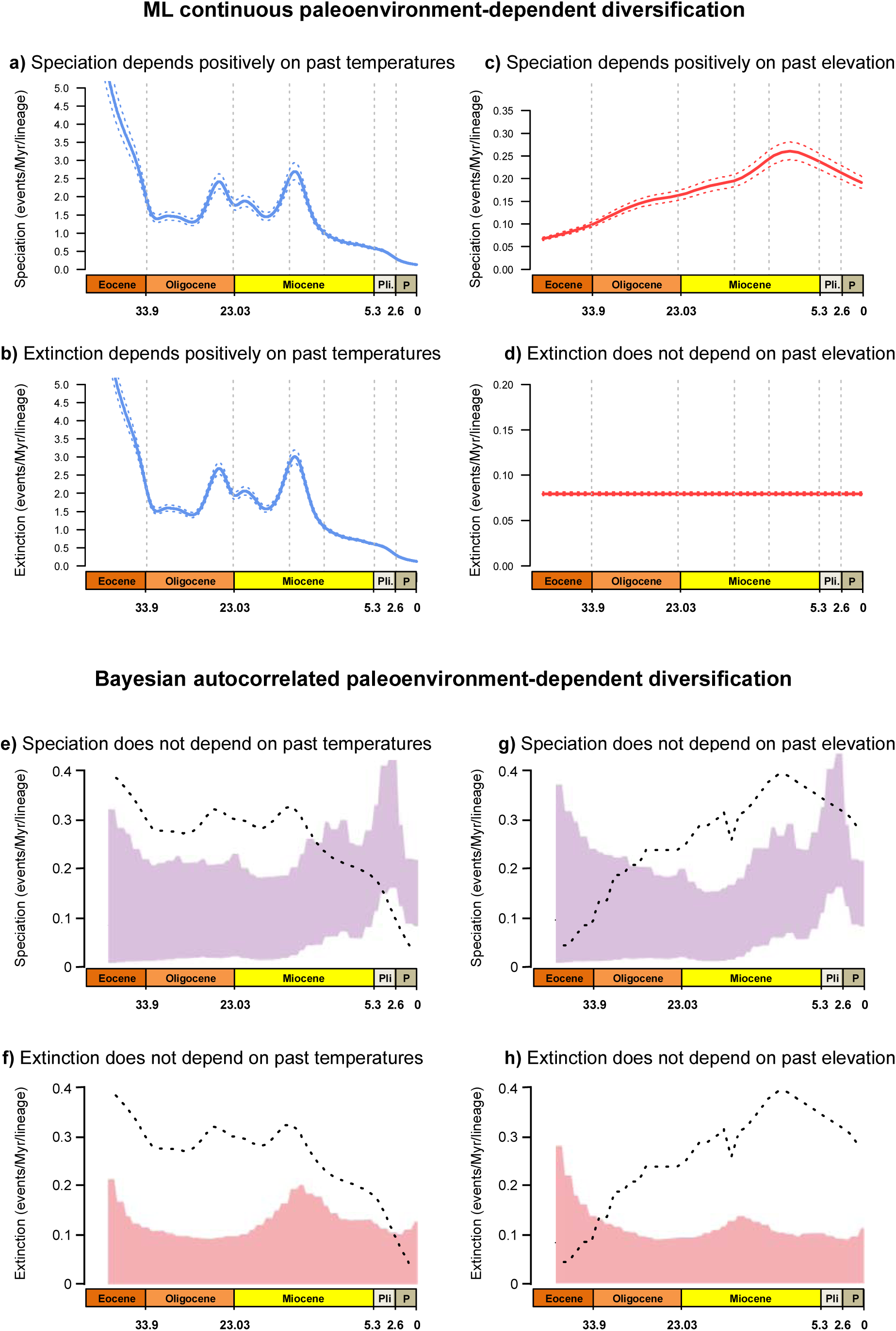
Paleoenvironment-dependent diversification processes in Apollo butterflies. ML continuous paleoenvironment-dependent models implemented in RPANDA (a-d) showedpositive dependence between paleotemperatures and speciation/extinction; the paleo-elevation-dependent diversification models did not fit the data better than the constant-rate or time-variable models. Bayesian inference of environment-dependent diversification with autocorrelated rates in RevBayes (e-h) supports a model in which both speciation and extinction are not dependent on temperature and elevation changes (represented by dotted lines, temperature and elevation values are given in Figure 2). Colored areas in speciation and extinction plots indicate the credibility interval, with the continuous line representing the median rate. Pli, Pliocene; and P, Pleistocene.

## Discussion

Seeking the determinants of species richness is a fundamental aspect of biology (Morlon 2014; Benton 2015). However, studies have generally focused on a single factor to explain a given diversity pattern, whether by historical events promoting changes in diversification and extinction (Barnosky 2001; Erwin 2009; Mayhew et al. 2008), or species ecology and clade competitions as the dominant drivers of diversification (Rabosky 2013; Silvestro et al. 2015). The combined influence of various factors has been tested less often (Drummond et al. 2012a; Bouchenak-Khelladi et al. 2015; Lagomarsino et al. 2016), and at a lower geographic and temporal scale or with a reduced set of methods. Further, these methods have mainly been implemented within a ML framework. In this study, we used existing and some novel methods under both ML and BI to evaluate potential factors driving diversification in Parnassiinae: some are related to a RQ mechanistic model of evolution, others to a CJ hypothesis.

### Coupled effect of mountain building and climate change

A long-held tenet in biology is that environmental stability leads to greater species richness than a changing environment, suggesting that species interactions are a dominant force (Wallace 1878). A recent debate has emerged between those arguing that physically dynamic ecosystems play a major role in diversification (Hoorn et al. 2013; Condamine et al. 2015a) and even promote adaptive radiation (Tan et al. 2013), and those maintaining that landscape changes have not been important in diversification history, instead advocating the role of large-scale dispersal events as drivers of speciation (Smith et al. 2014). In the Northern Hemisphere, tectonic events such as the closure of Turgai Sea or the building of Alpine and Himalayan orogenies have caused large-scale landscape and climatic changes (Sanmartín et al. 2001). As a result many new habitats were formed that created potential ‘new ecological spaces’ and favored speciation (Xing and Ree 2017). Also, the newly created corridors stimulated species exchange between communities, while also acting as barriers that separated formerly continuous populations (Brikiatis 2014). Alternatively, landscape dynamics (i.e. unlike long-term stable environments) may have shattered former habitats, leading organisms to extinction. The rise of the HTP has often been invoked as providing important opportunities to diversify (Fjeldså et al. 2012; Price et al. 2014; Wen et al. 2014; Favre et al. 2015; Hughes and Atchison 2015; Renner 2016), but its role as a driver has rarely been formally tested with diversification models (Xing and Ree 2017).

Our analyses detected higher *in-situ* speciation rates for mountain species than their lowland counterparts (with similar extinction rates), resulting in higher net diversification rates for mountain species. Yet, this pattern was not robust to phylogenetic shape artifacts (Rabosky and Goldberg 2015). Moreover, ML and BI-based paleo-elevation models found that an explicit link between diversification rates and periods of HTP uplift could be rejected in favor of a constant birth-death. These findings contrast with those of high-elevation taxa in the Andes reported to have diversified faster due to the rise of the Andes as compared to their low-elevation relatives (Lagomarsino et al. 2016; Pérez-Escobar et al. 2017). In contrast, clade rate-heterogeneous models (BAMM, RevBayes) and biogeographic analyses supported a link between diversification rates and the HTP orogeny within Parnassiini, particularly in *Parnassius*. A significant upshift in speciation rates was detected along the stem of *Parnassius*, coincident with the colonization of the HTP, while DEC inferred several speciation/vicariance events associated with the HTP orogeny (**Figure 4**). The latter is corroborated by the relatively high rate of allopatric (lowland/highland) speciation estimated by GeoSSE. These different results can partially be explained by the HTP rise providing novel, high-altitude habitats for colonization in *Parnassius*, and thereby promoting ecological divergence and initial rapid diversification (followed later by allopatric speciation). Yet, diversification was not strictly parallel to periods of mountain building. Conversely, the rise of the HTP – and concomitant climatic changes such as regional cooling and aridification – led to higher extinction rates in non-mountain-adapted Parnassiinae lineages, such as Luehdorfiini and Zerynthiini that survived west and east of the Himalayan range; several events of extinction are inferred within these lineages around the mid-Miocene. This mixed effect is probably responsible for the lack of a significant association between orogeny and speciation in tree-wide models, which is otherwise detected by clade-heterogeneous models.

Paleoclimate change is another abiotic factor postulated to be a major trigger of diversification in terrestrial organisms (Erwin 2009). CoMET, TreePar, and the ML-based paleotemperature-dependent model inferred a pattern of increased turnover (background extinction) rates associated with periods of global climate warming, such as the mid-Miocene Climate Optimum (MMCO, 15-17 Ma, Böhme 2003) or the late Oligocene Warming Event (LOWE, ~25 Ma, Zachos et al. 2008, **Figure 2**). DEC also reconstructed several events of (geographic) extinction in Central Asia and India along the stem branches subtending the crown diversification of extant genera in Luehdorfiini and Zerynthiini (**Figure 4**). These long branches are consistent with periods of high extinction rates related to the LOWE, which were eventually followed by mid-Miocene radiations, especially in *Parnassius* (the latter probably linked to the HTP rise). Paleotemperature and the HTP orogeny probably had a joint influence in the diversification of Parnassiinae. Mountain building is known to be responsible for regional climate change (Sepulchre et al. 2006; Armijo et al. 2015), and for changes in biodiversity patterns, both directly as a geographic barrier and indirectly through climate change (Hoorn et al. 2013). The HTP acted as a barrier to the influence of the Asian monsoons (Zhisheng et al. 2001) and induced considerable drying of Central Asia (Quade et al. 1989). In Parnassiinae, the HTP rise and subsequent periods of mountain building might have promoted diversification through the provision of “climate refugia”: locations where taxa survived periods of regionally adverse climate. Although usually associated with maintaining biodiversity through glacial–interglacial climate changes (Gavin et al. 2014), mountains can provide climate refugia during warming-aridification events (Migliore et al. 2013; Pokorny et al. 2015). Drastic range contractions within the non-mountain adapted Luehdorfiini and Zerynthiini (**Figure 4**) could have been driven by aridification events induced by HTP uplift and mid-Miocene climate warming, with *Hypermnestra* as a relict group that survived in the Zagros Mountains. In *Parnassius*, colonization of the HTP coincided with a boost of speciation, followed by dispersal to adjacent areas in the late Miocene-Pliocene. The long stem branches leading to extant Parnassiinae genera are likely the result of climate-driven extinction events associated with Cenozoic warming periods, such as the LOWE and the MMCO. During these periods, Parnassiinae species probably migrated toward cooler temperatures by colonizing the newly uplifted mountain ranges of the Zagros and HTP. This agrees with the climate refugia hypothesis, a pattern currently observed in Himalayan plants, which are shifting their distribution upward along with global warming (Padma 2014). We argue that the HTP acted as climate refugia during global warming events, in which pre-adapted parnassiines survived and diversified before re-colonizing ancestral geographic ranges. It would be interesting to test this hypothesis in other extra-Himalayan groups that diversified during the HTP orogeny (Favre et al. 2015), as well as in other mountains around the world for which paleo-elevation data is available (Fjeldså et al. 2012; Hoorn et al. 2013; Hughes and Atchison 2015).

### Ecological interactions via insect-plant diversification and host-plant shifts

Biological interactions among distantly related lineages probably played a major role in the diversification of clades over geological time (Van Valen 1973; Liow et al. 2015; Silvestro et al. 2015; Voje et al. 2015). Our analyses identified significant differences in speciation rates associated with different host-plants. Several models (BAMM, MuSSE, and Bayesian implementations of MuSSE and HiSSE in RevBayes) indicated that *Parnassius* subgenera associated with Crassulaceae-Saxifragaceae and Papaveraceae have significantly higher rates of speciation than their relatives feeding in other host plants (**Figures 5, 6**). Interestingly, the lowest speciation rates are associated with butterflies feeding on Aristolochiaceae, the inferred ancestral host plant of the Parnassiinae (Condamine et al. 2012). Our results concur with the ‘escape and radiate’ hypothesis of Ehrlich and Raven (1964), which predicts that the evolution of a herbivore insect trait (i.e. new tolerance of plant defenses) that allows colonizing/feeding on a novel host plant lineage would lead to a burst of diversification. The highly unbalanced extant species richness between Parnassiini and the clade Luehdorfiini+Zerynthiini, explained above by geographic and climate drivers, might also be explained by differential speciation rates related to their feeding habit, as well as the overall low ability of Parnassiinae for shifting between host plants (i.e. transition rates estimated MuSSE are close to zero).

Testing this hypothesis would require reconstruction of the diversification history of the host-plant clades; this would allow us, for example, to identify shifts that led to reciprocal changes in diversification in the host plants and to test whether these shifts were accompanied by matching ancestral geographic distributions between herbivore insects and plants at key events (Cruaud et al. 2012). Unfortunately, so far there is no complete phylogeny for any of the large host families for Parnassiinae, including Aristolochiaceae (~500-600 spp.), Papaveraceae (~700-800 spp.), Crassulaceae (~1300-1500 spp.), and Saxifragaceae (~600-700 spp.). However, we found a positive correlation between species richness of these plant lineages and the speciation rates of their insect feeders (MuSSE: R^2^=0.96, BAMM: R^2^=0.87; **Appendix 22** on Dryad). This suggests that fast-diversifying clades are associated with families offering a higher number of potential hosts (akin to niche availability). Janz et al. (2006) found a similar correlation between host-plant diversity and speciation rates in the Nymphalidae; they posited that recurring oscillations between host-plant expansions (i.e. incorporation of new plants into the feeding repertoire) and host specialization acted as a major driving force behind the diversification of plant-feeding insects.

In adaptive radiations, the expectation is an initial rapid burst of diversification, followed by a slowdown in speciation rates as niches are filled up; this is attributed to a diversity-dependence effect (Phillimore and Price 2008; Etienne et al. 2012). A shift to a new ecological resource (e.g. a new host plant) can be an adaptive breakthrough (Ehrlich and Raven 1964), providing higher opportunities for speciation at the start of the radiation, which significantly decrease as species accumulate. Though we did not detect this diversity-dependence effect in Parnassiinae, a pattern of speciation decreasing with standing diversity (and constant extinction rates) was found in *Parnassius*, supporting the hypothesis of this genus as an example of an adaptive radiation (Rebourg et al. 2006), driven by the evolution of a new host-plant association. The inference of similar extinction rates in clades of Parnassiinae feeding on different host plants does agree with Van Valen’s (1973) RQ hypothesis, which assumes constant extinction probabilities shared by all members of any given higher taxon.

### Interplay between the Red Queen and the Court Jester

Both abiotic and biotic factors drive species diversification, and biological radiations require both extrinsic conditions and intrinsic traits – acting in unison or sequentially over time – for success (Donoghue and Sanderson 2015). A lack of suitable methods and data has resulted in few studies attempting to merge both types of factors to explain evolutionary radiations (Bouchenak-Khelladi et al. 2015; Liow et al. 2015; Lagomarsino et al. 2016). Here, we argue that RQ and CJ-type factors are intimately linked, and they both promote and prevent species diversification. No single but multiple factors explain the diversification of Apollo butterflies, and the effect of these factors differed across clades. Past changes in climate and elevation during active orogenic periods jointly mediated the rapid radiation of mountain-dwelling *Parnassius*, but also triggered extinction events in non-mountain adapted clades Luehdorfiini and Zerynthiini. The ancestor of Luehdorfiini and Zerynthiini was reconstructed in the Oligocene as occupying a broad distribution range extending from the Caucasus region, the Iranian Plateau and the Zagros Mountains, through Central Asia to India (*red box* in **Figure 4**). Extinction in Central Asia and India around the Oligocene-Miocene, followed by independent eastward/northward colonizations by Luehdorfiini and Zerynthiini, could explain the current disjunct distribution pattern in these two tribes.

Our study also demonstrates that ecological traits and biotic interactions such as host-plant associations and clade-specific diversity-dependence may confer differential diversification capacity in closely related species co-occurring in the same rapidly changing environment. In particular, the radiation of *Parnassius* in the HTP seems to have been fostered by the joint effect of climate and geological changes associated with new biotic interactions. Mountain uplift promoted speciation by allopatric speciation, but also increased habitat heterogeneity, and led ultimately to the formation of new niches (ecological divergence). The colonization of these niches led to new associations with mountain host species, and potentially to the evolution of new morphological adaptive traits, e.g. denser setae on the body or altitudinal melanism. Although it is difficult to tease apart the role of key innovations (new traits that evolved allowing the invasion of a new niche) on ecological opportunity (the formation of a new environment that confers selective advantage fitness for the traits), a combination of extrinsic and intrinsic factors seems to have been necessary for the success of the *Parnassius* radiation. Parnassiinae species feed mostly on plant lineages of small size whose leaves grow close to the ground and which are characteristic elements of modern temperate vegetation: *Aristolochia* and *Asarum* (Aristolochiaceae) for Luehdorfiini and Zerynthiini, *Saxifraga* and *Sedum* (Saxifragaceae) and *Rhodiola* (Crassulaceae) for the subgenus *Parnassius*, or *Corydalis* (Papaveraceae) for the other subgenera of *Parnassius*. Zhang et al. (2014) inferred an origin of *Rhodiola* (70 spp., mid-Miocene; 12.1 Ma, 6.3-20.2 Ma) and *Sedum* (420 spp., late Oligocene) in the HTP, later spreading to the adjacent regions. Similarly, Ebersbach et al. (2017) placed the colonization of the HTP by genus *Saxifraga* (370 spp.) in the late Oligocene (20-34 Ma). Therefore, there was a synchronous temporal and geographic origin of *Parnassius* and its associated host plants in the HTP region. Conversely, Luehdorfiini and Zerynthiini were unable to evolve new traits or establish novel host-plant associations to invade the mountain and/or host-plant niches. This calls for further studies to decipher the genomic baseline for the adaptations to elevation and host plants by comparing Parnassiini and Luehdorfiini/Zerynthiini (Edger et al. 2015).

### Underlying model and inference framework in diversification analyses

Models of diversification have traditionally relied on time-continuous (independent) or autocorrelated rates in the way they model rate-shifts, with few studies applying both types for macroevolutionary inferences. Here we used both, in particular to estimate speciation and extinction rates depending on an environmental variable that itself varies through time. The first model is an ML implementation with a time-continuous birth-death model (Condamine et al. 2013). For the purpose of this study we implemented Bayesian paleoenvironment-dependent models but with autocorrelated (RevBayes) and time-continuous (TESS) rates of diversification. Our comparison across methods has revealed new aspects on the behavior of the models. The three methods are congruent on the role of orogeny (no effect of orogeny on diversification *per se*), but they disagree on the effect of temperature (**Figure 8**). The time-continuous models (RPANDA and TESS) support a positive dependence on speciation and extinction, while the autocorrelated model (RevBayes) shows no dependence. The results suggest that the lack of correlation found by the RevBayes paleoenvironmental model is directly a consequence of the assumption of autocorrelation in rates across time intervals. Thus, the difference lies in the underlying diversification model (autocorrelated vs. time-continuous rates) and not due to the inference framework (Bayesian vs. ML).

Our study shows that multiple approaches must be combined to fully address the diversification of clades in relation to abiotic and biotic factors. Rather than a single factor, the joint effect of multiple factors (biogeography, species traits, environmental drivers, and mass extinction) is responsible for current diversity patterns, and we recommend a correspondingly multifaceted strategy for the study of these patterns.

## Author Contributions

F.L.C., F.A.H.S. and I.S. designed the study; F.L.C. and F.A.H.S collected the data; F.L.C. analyzed the data with the help of J.R. on SSE models, S.H. on Bayesian implementation of diversification models, and I.S. on biogeography; F.L.C. and I.S. wrote the paper with significant contributions from J.R., S.H., and F.A.H.S.

## Supplementary Material

Data available from the Dryad Digital Repository: http://dx.doi.org/10.5061/dryad.[NNNN].

## Funding

This study has been funded by a Marie Curie Action (EU 7^th^Framework Programme) (BIOMME project, IOF-627684) to F.L.C. (jointly supervised by F.A.H.S. and I.S.), by a Natural Sciences and Engineering Research Council of Canada (NSERC), Discovery Grant to F.A.H.S., and by MINECO/FEDER (CGL2015-67489-P) to I.S.

## Acknowledgments

We thank Sylvain Piry for help on the script for GenBank, Mark Miller for assistance on the CIPRES cluster.

## APPENDICES

**Appendix 1.** All sequence data used for this study (a file is generated per gene).

**Appendix 2.** The individual gene alignments as recovered by MAFFT. Results of the Bayesian phylogenetic analyses for each gene, and an explanation of the results.

**Appendix 3.** The total-evidence matrix (including molecular and morphological data) used for the phylogenetic placement of fossils with MrBayes.

**Appendix 4.** The BEAST files for the Bayesian dating analyses (the tree prior can be a Yule process or a birth-death model, and the dataset can include or not the morphological data). **[H2]Appendix 5.** The current geographic species distribution data of all Parnassiinae as coded present (1) or absent (0) in all ten geographic areas (Western Palearctic, North Africa, Turkey, Central Asia, Himalaya, India, Mongolia, Siberia, China-Japan, and Western Nearctic).

**Appendix 6.** The time-stratified biogeographic model used for DEC analyses (time slices represent geological epochs or stages in the Cenozoic).

**Appendix 7.** Paleo-elevation for the Himalayan and Tibetan compiled from the literature.

**Appendix 8.** Description of the Bayesian episodic environment-dependent birth-death model.

**Appendix 9.** Results of PartitionFinder performed on the concatenated molecular dataset.

**Appendix 10.** Time-calibrated trees of Parnassiinae as estimated by BEAST following four different analyses.

**Appendix 11.** Results of the model comparison for the dating analyses based on marginal likelihood estimates and Bayes factors.

**Appendix 12.** Biogeographic history of Parnassiinae as estimated by DEC.

**Appendix 13.** Results from the diversity-dependence diversification analyses in DDD.

**Appendix 14.** Results of the MuSSE and GeoSSE analyses performed on 200 trees randomly taken from the Bayesian dating analysis. Models are ranked by AICc.

**Appendix 15.** Plot of the difference between speciation rates between all traits. When the difference overlaps zero (vertical red bar), the speciation rates are not significantly different.

**Appendix 16.** Robustness of the SSE models with simulation tests, HiSSE analyses and an implementation in RevBayes. For the simulation, the difference of fit between the best model and the reference model is shown with the red vertical line for real data, and in black for simulated data. HiSSE and RevBayes agree with the MuSSE models on host plants.

**Appendix 17.** Summary of diversification models in BAMM compared across a gradient of values for the Poison process governing the number of rate shifts.

**Appendix 18.** Credible set of configuration shifts inferred with BAMM and five different values of the Poison prior. It shows the distinct shift configurations with the highest posterior probability. For each shift configuration, the locations of rate shifts are shown as black circles, with circle size proportional to the marginal probability of the shift.

**Appendix 19.** Rate-through-time plot as inferred with RevBayes for Parnassiinae. Net diversification rates significantly changed and increased along the stem of the genus *Parnassius*, in agreement with the rates as estimated with BAMM.

**Appendix 20.** Rate-through-time plot as inferred with CoMET for Parnassiinae. The analyses detected one possible mass extinction around 15 Ma and one speciation rate shift around 3.5 Ma, in agreement with two TreePar analyses allowing or not the mass extinction.

**Appendix 21.** Credibility intervals of the correlation parameters for the Bayesian (RevBayes) environment-dependent diversification models.

**Appendix 22.** Correlation (linear regression) between speciation rates as inferred with MuSSE (a) and BAMM (b) and the species richness of host plants on which each parnassiine clade is feeding. In both cases, a strong and positive correlation is found.

